# Molecular characterization of nodose ganglia development reveals a novel population of Phox2b+ glial progenitors in mice

**DOI:** 10.1101/2023.07.25.550402

**Authors:** Elijah D. Lowenstein, Aristotelis Misios, Sven Buchert, Pierre-Louis Ruffault

**Author notes:** Corresponding author. Present address: Department of Medicine, Division of Endocrinology, Diabetes and Metabolism. Beth Israel Deaconess Medical Center and Harvard Medical School, Boston, MA 02115, USA. Electronic address.

## Abstract

The vagal ganglia, comprised of the superior (jugular) and inferior (nodose) ganglia of the vagus nerve, receive somatosensory information from the head and neck, or viscerosensory information from the inner organs, respectively. Developmentally, the cranial neural crest gives rise to all vagal glial cells and to neurons of the jugular ganglia, while the epibranchial placode gives rise to neurons of the nodose ganglia. Crest-derived nodose glial progenitors can additionally generate autonomic neurons in the peripheral nervous system, but how these progenitors generate neurons is unknown. Here, we found that some Sox10+ neural crest-derived cells in, and surrounding, the nodose ganglion transiently expressed Phox2b, a master regulator of autonomic nervous system development, during early embryonic life. Our genetic lineage tracing analysis revealed that despite their common developmental origin and extreme spatial proximity a substantial proportion of glial cells in the nodose, but not in the neighboring jugular ganglia, have a history of Phox2b expression. Lastly, we used single cell RNA-sequencing (scRNA-seq) to demonstrate that these progenitors give rise to all major glial subtypes in the nodose ganglia, including Schwann cells, satellite glia and glial precursors, and mapped their spatial distribution by *in situ* hybridization. Our work demonstrates that these crest-derived nodose glial progenitors transiently express Phox2b, give rise to the entire complement of nodose glial cells and display a transcriptional program that may underlie their bipotent nature.

**Significance statement:** The nodose ganglia contain sensory neurons that innervate many inner organs and play key roles in homeostatic behaviors such as digestion, regulation of blood pressure and heart rate, and breathing. Nodose sensory neurons are supported by nodose glial cells, which are understudied compared to their neuronal neighbors. Specifically, the genetic program governing their development is not fully understood. Here, we uncover a transcriptional program unique to nodose glial cells (transient expression of Phox2b) that resolves the 40-year-old finding that nodose glial progenitors can also give rise to autonomic neurons (whose development depends on Phox2b expression). Lastly, we leveraged single cell RNA-sequencing to identify the four major subtypes of nodose glial cells and used subtype specific marker genes to map their spatial distribution.

## Introduction

Viscerosensory neurons in the nodose ganglia are important to maintain bodily homeostasis and reside intermingled with resident peripheral glial cells (Chang et al., 2015; Zeng et al., 2018; Vermeiren et al., 2020; Prescott and Liberles, 2022; Lowenstein et al., 2023). Peripheral glial cells, including both Schwann cells and satellite glia, regulate synaptic connectivity, provide nutrients and structural support, as well as myelinate neurons in the peripheral nervous system (Riethmacher et al., 1997; Véga et al., 2003; Hanani, 2005; Fields and Burnstock, 2006; Jessen et al., 2015; Monk et al., 2015; Avraham et al., 2020). Single cell RNA-sequencing (scRNA-seq) has begun to elucidate the diversity of peripheral glial cells in numerous peripheral nerves as well as in autonomic and somatosensory ganglia, revealing transcriptomic differences in glial cell types across the peripheral nervous system (Wolbert et al., 2020; Gerber et al., 2021; Tasdemir-Yilmaz et al., 2021; Avraham et al., 2022; Mapps et al., 2022). However, the number of glial types in the viscerosensory nodose ganglia and their transcriptomic profiles that support the vast array of nodose neuron types is not known.

Developmentally, cells of the nodose ganglia arise from two neurogenic niches: glial cells derive from the cranial neural crest while neurons derive from the epibranchial placode (Vermeiren et al., 2020). The differential contributions of the crest and placode were elucidated by a series of seminal quail-chick embryo transplantation studies where isotropic and isochronic grafts of quail crest or placode were transplanted into a chick host (Narayanan and Narayanan, 1980; Ayer-Le Lievre and Le Douarin, 1982; D’amico-Martel and Noden, 1983). These chimeras demonstrated that crest-derived non-neuronal nodose cells could generate autonomic neurons when back-transplanted into the neural crest of a younger host (Ayer-Le Lievre and Le Douarin, 1982; Le Douarin, 1986; Fontaine- Perus et al., 1988; Douarin et al., 1991). Although crest derivatives only generate glial cells in the nodose ganglia, they may acquire other cell fates when placed in a new permissive environment.

In mice the viscerosensory nodose ganglia, i.e. inferior ganglia of the vagus nerve, are directly abutting the somatosensory jugular ganglia, i.e. superior ganglia of the vagus nerve. In contrast to the nodose ganglia, both neurons and glial cells of the jugular ganglia derive from the cranial neural crest (Vermeiren et al., 2020). Thus, the same neurogenic niche generates both jugular and nodose glial cells, but it is currently unknown whether there are molecular differences between nodose and jugular glial cell development.

With the advent of molecular biology and the development of genetic lineage tracing, the broad transcriptional programs necessary for peripheral glial cell and neuron development were revealed. All peripheral glial cells derive from the neural crest, an embryonic niche that depends on the Wnt signaling pathway (Jessen and Mirsky, 2005; Jacob, 2015; Erickson et al., 2023). The successful differentiation of neural crest cells into the full complement of peripheral glial cells relies on the transcription factor *Sox10*, whose ablation precludes the generation of Schwann and satellite glial cells (Kuhlbrodt et al., 1998; Britsch et al., 2001). In contrast, neurons of the autonomic nervous system, including sympathetic, parasympathetic, enteric and cranial sensory neurons, depend on the related homeobox transcription factors *Phox2a* and *Phox2b* for their development and maintenance (Pattyn et al., 1997, 1999; Yang et al., 2002). Thus, transgenic mouse strains expressing recombinases in cells expressing *Wnt1* and *Phox2b* provide tools to study crest and placode derivatives, respectively.

Despite that fact that Phox2a/b instruct neuronal fates, in parasympathetic preganglionic nerves, some nerve-associated Sox10+ glial precursors transiently express Phox2b before adopting a mature Schwann cell identity (Dyachuk et al., 2014; Espinosa-Medina et al., 2014). Recent studies used scRNA-seq to dissect the transcriptional changes coinciding with cell fate acquisition and found that many cells initially express competing gene modules (e.g. neuronal and glial), before selecting one module over the other (e.g. neuronal > glial) (Soldatov et al., 2019; Kastriti et al., 2022). However, whether the bipotent nature of nodose glial cells revealed by the early quail-chick transplantation experiments is reflected in their transcriptional program during development remains to be investigated. Specifically, we hypothesize that one should be able to find nodose glial cells that transiently express Phox2b, the master transcription factor that is required for proper autonomic neuron development.

Here, we used genetic lineage tracing and scRNA-seq to identify the transcriptional programs leading to the development of neurons and glial cells in the nodose ganglia. We show that a significant proportion of nodose, but not jugular, glial cells derive from a subset of Wnt1*+* neural crest progenitors that transiently express Phox2b during early embryogenesis. Our scRNA-seq results revealed the developmental potential of neural crest cells that have a history of Phox2b expression to include all major glial subtypes, i.e. myelinating (MSC) and non-myelinating Schwann cells (NMSC), but also satellite glia (SG) and glial precursors (GP). These results suggest that the bipotent character of nodose glial cell progenitors is reflected in their transcriptional program and differentiates them from jugular glial cell progenitors despite their shared developmental origins and spatial proximity.

## Materials and methods

### Mouse lines

All experiments were conducted according to regulations established by the Max Delbrück Center for Molecular Medicine, LAGeSo (Landesamt für Gesundheit und Soziales). *Ai14* (#007908) mice were obtained from the Jackson Laboratory (Madisen et al., 2010). *Tau^nLacZ^* (Hippenmeyer et al., 2005) mice were provided by Silvia Arber (Biozentrum, University of Basel, Switzerland). *Wnt1^Cre^* (Danielian et al., 1998) mice were provided by Andrew McMahon (USC, Los Angeles, USA). *Phox2b^Cre^* (D’Autréaux et al., 2011), and *Phox2b^FlpO^* (Hirsch et al., 2013) mice were provided by Jean-François Brunet (Institut de Biologie de l’ENS, Paris, France). *R26^ds-nGFP^*(Britz et al., 2015) mice were provided by Martyn Goulding (Salk Institute, San Diego, USA). *Gt(ROSA)26Sor<tm2.1Sia>* mice were provided by Shinichi Aizawa (RIKEN Center for Developmental Biology, Kobe, Japan); we refer to them as *R26^nGFP^* mice, as they express a nuclear GFP upon cre-mediated stop cassette excision (Abe et al., 2011). Mice were housed at room temperature (23°C), humidity (56%), and with a 12-hour light-dark cycle.

### Tissue processing

Pregnant dams were staged using the presence of a vaginal plug. The day that a vaginal plug was visible was designated as embryonic (E) day 0.5. Embryos were delivered by cesarean section and fixed for 1-3 hours in 4% PFA in PBS at 4°C. They were washed for 2 x 10 minutes in PBS at room temperature before cryopreservation overnight at 4°C in 15% sucrose in PBS, and then overnight at 4°C in 30% sucrose in PBS. Cryopreserved embryos were washed once and embedded in Tissue-Tek OCT (Sakura) in a plastic block before being stored at -80°C. Embryos were sectioned at 20 µm using a cryostat. Vagal ganglia were dissected from postnatal (P) day 4 pups and fixed for 1 hour in 4% PFA in PBS, before cryopreservation for 30 minutes in 15% sucrose in PBS followed by 30 minutes in 30% sucrose in PBS all at 4°C. Vagal ganglia were washed once in Tissue-Tek OCT (Sakura) and embedded in Tissue-Tek OCT (Sakura) in a plastic block before being stored at -80°C. Vagal ganglia were sectioned at 16 µm using a cryostat. After sectioning slides were stored at -80°C until further use.

### Immunofluorescence

Immunofluorescence experiments were performed on cryosections using the following primary antibodies: rat anti-GFP (Nacalai Tesque, GF090R, 1:1000), goat anti-Phox2b (R&D Systems, AF4940, 1:200), rabbit anti-Sox10 (Abcam, ab227680, 1:200), goat anti-Sox10 (R&D Systems, AF2864, 1:200), guinea pig anti-Tlx3 (1:5000, kindly provided by Thomas Müller), goat anti-TrkA (R&D Systems, AF175, 1:500), goat anti-TrkB (R&D Systems, AF1494, 1:300), chicken anti-βgal (Abcam, ab134435, 1:1000), chicken anti-GFP (Aves Labs, GFP-1020, 1:500), rabbit anti-RFP (Rockland, 600-401-379-RTU, 1:500), goat anti-RFP (Biorbyt, orb334992, 1:500), and rabbit anti-Phox2b (1:500, kindly provided by Jean-François Brunet). Slides were removed from -80°C and thawed at 37°C for 30 minutes. Slides were washed once in PBS for 1 minute and once in PBX (PBS with 0.2% Triton X-100) for 3 minutes at room temperature. Sections were then incubated in blocking solution (PBX with 5% horse serum) for 1 hour at room temperature. Primary antibodies were diluted in blocking solution and left on the sections overnight at room temperature. Slides were then washed twice in PBS for 30 seconds, twice in PBS for 10 minutes and once in PBX for 10 minutes at room temperature. Species specific secondary antibodies coupled to Cy2-, Cy3- or Cy5 (Jackson ImmunoResearch, 1:500) and DAPI were diluted in blocking solution and left on the sections for 1 hour at room temperature. Slides were washed three times in PBS for 10 minutes at room temperature before mounting them with Immu-Mount (Thermo Fisher Scientific).

### Vagal ganglia dissociation, neuronal depletion and cell sorting

Vagal ganglia were dissected from 4 *Phox2b^Cre^;Ai14* mice at P4 of either sex and placed into ice cold Ringer’s solution (0.125 M sodium chloride, 1.5 mM calcium chloride dihydrate, 5 mM potassium chloride, 0.8 mM sodium phosphate dibasic). Ganglia were transferred to a 1.5 ml Eppendorf tube with 1.5 ml HBSS without calcium, magnesium and phenol red (Thermo Fisher Scientific), 3 µl saturated NaHCO_3_, 1 mg L-Cysteine and 60 U Papain (Sigma-Aldrich), placed at 37°C and shaken gently for 12 minutes. The 1.5 ml Eppendorf tube was centrifuged at 300 RCF and the solution was discarded, taking care not to disturb the ganglia. A prewarmed (37°C) enzyme solution containing 1 ml HBSS without calcium, magnesium and phenol red (Thermo Fisher Scientific), 4 mg Collagenase NB4 (Serva) and 4.6 mg Dispase II (Gibco) was added to the ganglia, placed at 37°C and shaken gently for 12 minutes. The 1.5 ml Eppendorf tube was centrifuged at 300 RCF and the solution was discarded, taking care not to disturb the ganglia. Prewarmed (37°C) 0.5 ml HBSS without calcium, magnesium and phenol red (Thermo Fisher Scientific) was added to the ganglia, which were triturated with a 1 ml plastic pipette tip until the ganglia began to dissociate, and then with a 200 µl plastic pipette tip until the ganglia were completely dissociated. The dissociated ganglia were passed twice through a 40 µm filter (Falcon) and DAPI (Sigma) was added to a final concentration of 300 nM to label dead cells before sorting. We sorted tdTomato-positive/DAPI-negative cells into 4 96-well plates using an ARIA Sorter III (BD) and BD FACSDiva software 8.0.1.

### Library generation and sequencing

Single cell RNA-sequencing was done following the CEL-Seq2 protocol (Hashimshony et al., 2016). cDNA Libraries were prepared for 4 96-well plates, pooled together, and sequenced on 1 lane of a HiSeq 1500 (Ilumina) by the next generation sequencing core facility at the Max Delbrück Center for Molecular Medicine.

The sequencing dataset generated in this study has been deposited in the GEO repository (GEO accession number GSE237947) and are publicly available.

### Analysis of the P4 vagal ganglia CEL-Seq2 data

Data processing and gene quantification was performed using dropseq-tools v2.0 and picard-tools v2.18.17. We used the standard pipeline for Drop-seq data, with the necessary adaptations to analyze the CEL-Seq2 data. We removed the bead barcode correction steps, and in the DigitalExpression quantification we inputted the list of 96 CEL-Seq2 barcodes. Alignment was performed with STAR v2.5.3a (Dobin et al., 2013). We used the GRCm38 genome and the annotation from the GRCm38.p4 assembly. One cell barcode was missing from all plates, likely due to an error during library preparation, and one well had 10 times more reads and UMIs than the next highest well and was removed as an outlier. Two thresholds were set to filter out wells without cells or with damaged cells. We set a lower threshold of 5,000 UMIs (unique molecular identifiers) per cell, and a threshold of 10% of reads coming from mitochondrial genes. These thresholds filtered out 88 cells. After quality control we removed 88+4+1=93 cells (24% of the total cell count), leaving 291 cells for downstream analyses. Downstream analyses were performed with Seurat v3.0 (Butler et al., 2018). Seurat was run using the default parameters.

### Single molecule fluorescent *in situ* hybridization (RNAscope) and immunohistology

Single molecule *in situ* fluorescent hybridization (smFISH) was performed using the RNAscope Multiplex Fluorescent Reagent Kit V2 from ACDbio according to the manufacturer’s instructions. Briefly, 16 µm thick vagal ganglia sections were thawed at 37°C for 30 minutes and post-fixed in 4% PFA in PBS for 15 minutes before washing in PBS and continuing with the manufacturer’s instructions. For combining immunohistology with RNAscope we proceeded until the hydrogen peroxide wash, then washed the sections in PBS and incubated them at 4°C overnight with the primary antibody diluted in Co-Detection Antibody Diluent (obtained from ACDBio). The sections were washed in PBS and the protease treatment was performed using Protease III. The RNAscope protocol was then continued, the sections washed in PBS and incubated with the secondary antibody for 1 hour at room temperature in blocking solution (PBS with 0.2% Triton X-100 and 5% normal horse serum). Sections were washed in PBS and mounted with ProLong Gold Antifade mountant (ThermoFisher). We used the following probes in this study: Phox2b (407861-C2 and 407861-C3), Sox8 (454781-C2), Sox10 (435931), Scn7a (548561), Fabp7 (414651), Pou3f1 (436421-C2), and Ki67 (416771-C2).

### Microscopy, Image analysis and Cell quantifications

Photomicrographs were acquired using an LSM 700 laser scanning confocal microscope (Zeiss) with a 20x objective. Some images were acquired using the tile-scan mode with a 15% overlap between tiles and stitched using ZEN2012 software. Brightness and contrast were adjusted using ImageJ on the entire photomicrograph. Cell counts were performed by hand in ImageJ using the multi-point tool in a non-blind manner on non-consecutive sections. Between 2- 10 sections were counted per animal per experiment.

### Statistics

Statistics were calculated using Prism 6 (GraphPad). Data are plotted as scatter dot plots with the mean of each animal displayed as a dot, and the standard deviation shown. Statistical significance between group means was calculated either using a two-tailed t-test (two groups, Figure 3) or a one-way analysis of variance followed by Tukey’s post-hoc test (more than two groups, Figure 1).

**Figure 1.**
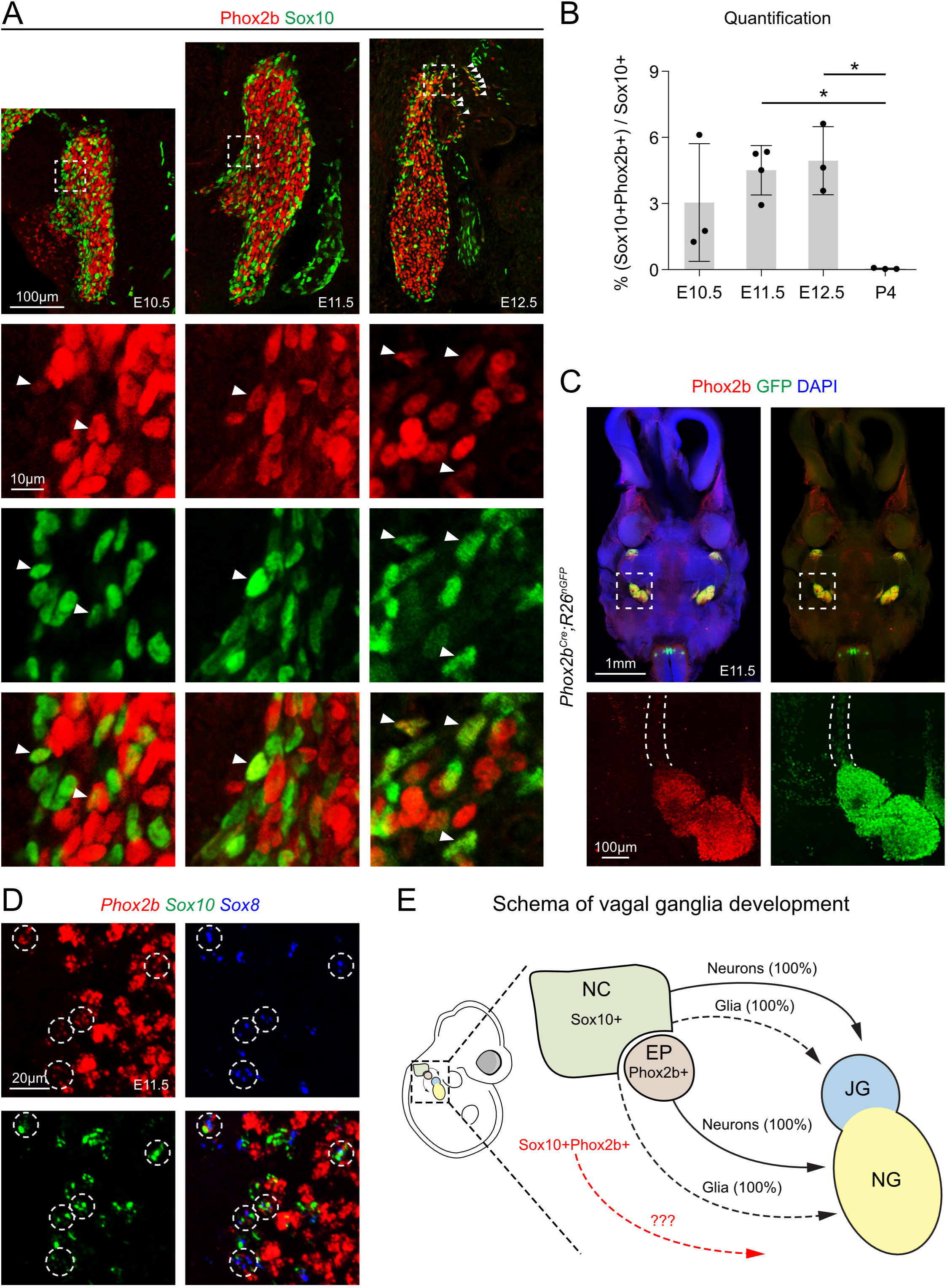
Some Sox10+ cells in the nodose anlage transiently express Phox2b during early development. **(A)** Immunohistology of sagittal sections of the nodose anlage at E10.5, E11.5 and E12.5 using antibodies against Phox2b (red) and Sox10 (green). Magnifications of the boxed regions are shown below. Arrows mark Sox10+Phox2b+ cells. **(B)** Quantifications revealed that 3.0±2.7, 4.5±1.1, 4.9±1.5 of Sox10+ cells in the nodose ganglia co-express Phox2b at E10.5, E11.5 and E12.5, respectively. Shortly after birth at P4 0.0±0.0% of Sox10+ cells express Phox2b and these proteins are exclusively present in nodose glial cells and neurons, respectively. N=3-4. **(C)** Immunohistology of a CUBIC cleared 300µm transverse section from an E11.5 *Phox2b^Cre^;R26^nGFP^*embryo using antibodies against Phox2b (red) and GFP (green). DAPI was used as a counterstain. One nodose ganglion is shown magnified below, and the vagus nerve is outlined with dotted lines. Note that there are many GFP+Phox2b- cells along the nerve. **(D)** smFISH (RNAscope) against *Phox2b* (red), *Sox10* (green) and *Sox8* (blue) mRNA. White dotted circles show *Phox2b*+*Sox10*+*Sox8*+ cells. **(E)** Schema showing vagal ganglia development. The neural crest (NC, olive green) contains Sox10+ progenitors that give rise to neurons (solid lines) and glial cells (dotted lines) of the jugular ganglia (JG, blue) and glia of the nodose ganglia (NG, yellow). NG neurons derive from the Phox2b+ epibranchial placode (EP, red). The derivatives from Sox10+Phox2b+ cells are unknown and are shown in red. Data are represented as mean ± SD, *p < 0.05, Ordinary one-way ANOVA with Tukey’s multiple comparisons test (B). Abbreviations: JG, jugular ganglia; NG, nodose ganglia; NC, neural crest; EP, epibranchial placode.

## Results

### A fraction of nodose ganglia cells co-express Sox10 and Phox2b during early embryonic development

Cells derived from the cranial neural crest (Wnt1+) and epibranchial placodes (Phox2a+) generate the glial cells and neurons of the nodose ganglia, respectively (Ayer-Le Lievre and Le Douarin, 1982; Morin et al., 1997; Pattyn et al., 1997; Dorsky et al., 1998). Interestingly, previous work showed that nodose glial cells retain the capacity to generate crest-derived autonomic neurons in early embryonic life (Ayer-Le Lievre and Le Douarin, 1982; Le Douarin, 1986; Fontaine-Perus et al., 1988; Douarin et al., 1991). Given that Phox2b is necessary for the development and survival of autonomic neurons (Pattyn et al., 1999), we first asked whether nodose neural crest progenitors express Phox2b during development. We used immunohistology to examine the expression of Sox10 and Phox2b, two transcription factors that respectively mark glial cells and neurons in the nodose ganglion between embryonic (E) day 10.5-12.5 (Figure 1A). At E10.5-E11.5, most Phox2b+ neurons have delaminated from the epibranchial placode and begun to coalesce in the nodose ganglia alongside Sox10+ glial cells derived from the neural crest and from nerve-associated crest progenitors (Fode et al., 1998; Zou et al., 2004). We found that around 5% of Sox10+ cells in the nodose anlage co-expressed Phox2b between E10.5-12.5. The presence of Sox10+Phox2b+ cells was no longer observed by postnatal (P) day 4 (quantified in Figure 1B). To exclude that these early Sox10+Phox2b+ cells represent an immature state of nodose neurons, we co-stained nodose ganglia with antibodies against Sox10 and the early post-mitotic neuron marker Tlx3 at E11.5 and E12.5 (Kondo et al., 2008). This revealed that Sox10+ cells co-expressing Tlx3 were extremely rare (1/1911 cells at E11.5, n=3; 1/3047 cells at E12.5, n=3), suggesting that Sox10+ cells are not immature neurons (Extended Data Figure 1-1). Thus, cells co-expressing Sox10 and Phox2b during early nodose ganglia development correspond to glial cells or nerve-associated crest progenitors.

To better visualize all cells with a history of Phox2b expression we designed a lineage tracing experiment where we crossed a *Phox2b^Cre^* mouse line with the reporter allele *R26^nGFP^* that expresses nuclear GFP upon Cre-mediated recombination, and performed immunohistology against Phox2b and GFP. The resulting *Phox2b^Cre^;R26^nGFP^*mice were analyzed at E11.5 (Figure 1C). We observed many GFP+Phox2b- cells along the vagus nerve (outlined with a dotted white line in Figure 1C), suggesting that a subset of neural crest-derived cells transiently express Phox2b (i.e. expressed Phox2b prior to E11.5 and are thus GFP+, but already down-regulated Phox2b expression and are Phox2b-) on their way to the nodose ganglion.

Recent research indicated that nerve-associated crest cells, also known as Schwann cell precursors, generate neurons and glial cells in parasympathetic ganglia (Dyachuk et al., 2014; Espinosa-Medina et al., 2014). In particular, starting at E10.5 all neural crest cells begin to associate themselves with peripheral nerves or ganglia, and by E11.5 they are bona fide Schwann cell precursors defined by their expression of *Sox8*, a recently described marker for late migrating crest cells/Schwann cell precursors that begin to attach to peripheral nerves at E11.5 (Adameyko et al., 2012; Dyachuk et al., 2014; Espinosa-Medina et al., 2014; Furlan et al., 2017; Kastriti et al., 2022). To clarify whether the Sox10+Phox2b+ cells described above correspond to Schwann cell precursors, we performed single molecule fluorescent *in situ* hybridization (smFISH) at E11.5 using probes against *Sox10*, *Phox2b*, and *Sox8* mRNA (Figure 1D). This revealed that many *Sox10*+*Phox2b*+ cells were also positive for the Schwann cell precursor marker *Sox8* (white circles, Figure 1D). The fact that i) many GFP+Phox2b- cells in *Phox2b^Cre^;R26^nGFP^*mice were found along the vagus nerve together with ii) our *in situ* data showing that many *Sox10+Phox2b+* cells also co-express *Sox8*, suggests that they are Schwann cell precursors. A schema of vagal ganglia development is shown in Figure 1E, and the Sox10+Phox2b+ cells described here are highlighted in red.

### A large proportion of nodose glial cells derived from the neural crest have a history of Phox2b expression

To identify all nodose glial cells with a history of Phox2b expression, we carried out a long-term lineage tracing experiment using *Phox2b^Cre^*;*R26^nGFP^*animals and performed immunofluorescence at P4 (see Figure 2A for a schematic display of the genetic strategy). Nodose ganglia were identified by their expression of Phox2b, and distinguished from the neighboring Phox2b- jugular ganglia by their expression of TrkA (Figure 2B) (Nassenstein et al., 2010; Kupari et al., 2019). We found that in the nodose ganglia of *Phox2b^Cre^;R26^nGFP^*mice almost all Phox2b+ cells expressed GFP (97.3±2.8%, n=3), but surprisingly only half (44.1±11.1%, n=3) of GFP+ cells co-expressed Phox2b (Figure 2C, left). Next, we stained the nodose ganglia with antibodies against Sox10 and Tlx3 to determine the identity of GFP+Phox2b- cells in *Phox2b^Cre^;R26^nGFP^* animals. This indicated that a large proportion (50.4±8.3%, n=3) of GFP+ cells in the nodose ganglia expressed Sox10, whereas the remainder (53.4±6.8%, n=4) expressed Tlx3 (Figure 2C middle and right). Lastly, we also determined the proportion of Sox10+ cells that co-expressed GFP and found that roughly 35.2±2.3% (n=3) of all nodose glial cells have a history of Phox2b expression (Figure 2C middle). Thus, many nodose glial cells transiently expressed Phox2b during their development.

**Figure 2.**
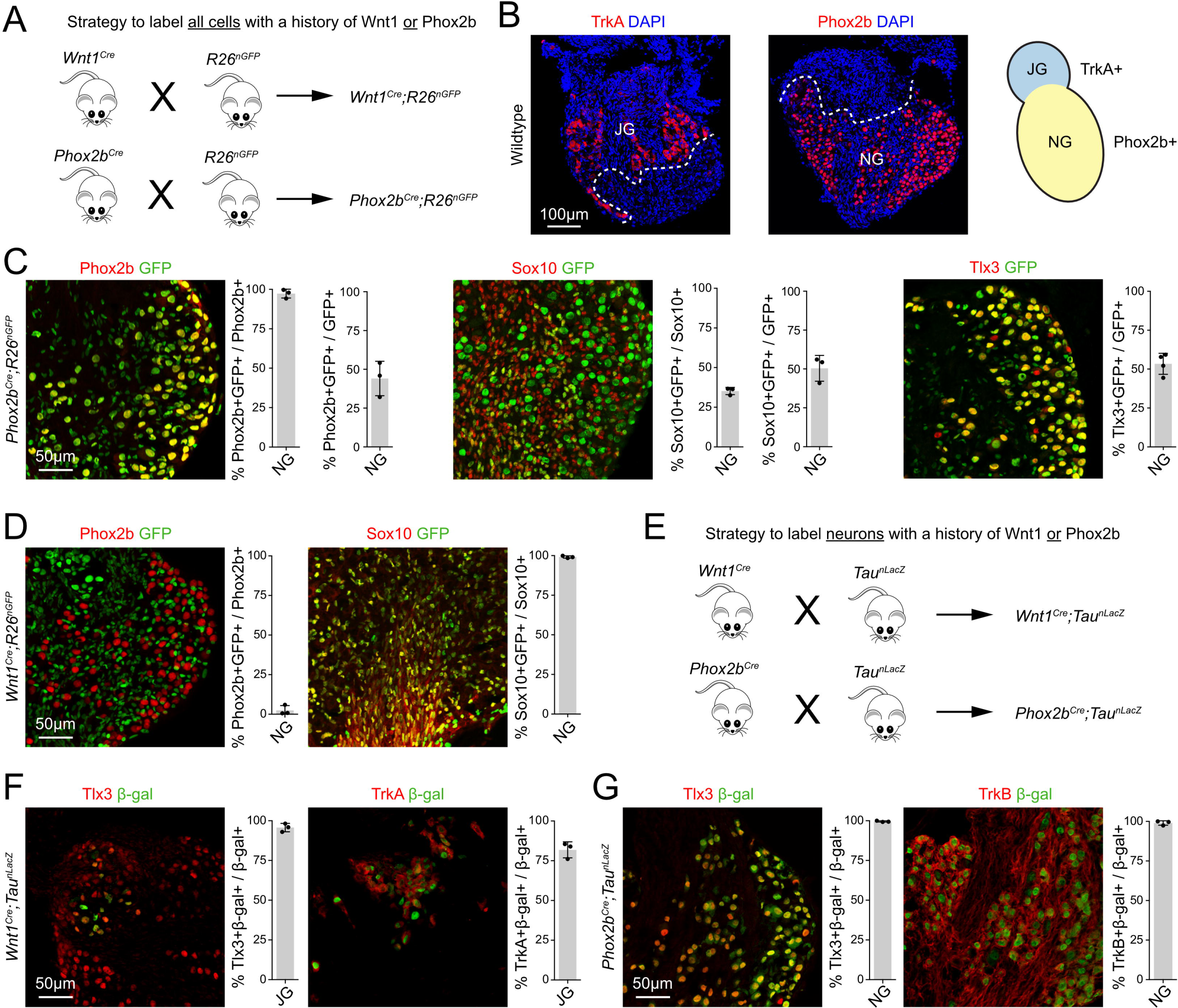
Nodose glial cells have a history of Phox2b expression. **(A)** Schema outlining the genetic lineage tracing strategy used to label all cells with a history of either Wnt1 or Phox2b. We crossed *Wnt1^Cre^* or *Phox2b^Cre^* animals with *R26^nGFP^* reporter mice to generate *Wnt1^Cre^;R26^nGFP^* and *Phox2b^Cre^;R26^nGFP^* mice. In these animals all cells that expressed or express Wnt1 (in *Wnt1^Cre^;R26^nGFP^*) or Phox2b (in *Phox2b^Cre^;R26^nGFP^*), will express a nuclear GFP. **(B)** We used immunohistology against TrkA and Phox2b to distinguish between the neighboring TrkA+Phox2b- jugular ganglion (JG) and the TrkA-Phox2b+ nodose ganglion (NG). Dotted white lines show the boundary between the jugular and nodose ganglia. For clarity a vagal ganglion is shown schematically on the right. **(C)** Immunohistology experiments in the nodose ganglia of *Phox2b^Cre^;R26^nGFP^* mice at P4. Immunohistology against GFP (green) and Phox2b (red, left), quantifications show that 97.3±2.8% of Phox2b+ cells are GFP+, but surprisingly only 44.1±11.1% of GFP+ cells are Phox2b+, n=3. Immunohistology against GFP (green) and Sox10 (red, middle), quantifications show that unexpectedly 35.2±2.3% of Sox10+ cells are GFP+, and 50.4±8.3% of GFP+ cells are Sox10+, n=3. Immunohistology against GFP (green) and Tlx3 (red, right), quantification shows that 53.4±6.8% of Tlx3+ cells are GFP+, n=4. **(D)** Immunohistology experiments in the nodose ganglia of *Wnt1^Cre^;R26^nGFP^* mice at P4. Immunohistology against GFP (green) and Phox2b (red, left), quantifications show that 2.4±3.0% of Phox2b+ cells are GFP+, n=3. Immunohistology against GFP (green) and Sox10 (red, middle), quantifications show that 99.1±0.7% of Sox10+ cells are GFP+, n=3. **(E)** Schema outlining the genetic lineage tracing strategy used to label neurons with a history of either Wnt1 or Phox2b. We crossed *Wnt1^Cre^* or *Phox2b^Cre^*animals with *Tau^nLacZ^* reporter mice to generate *Wnt1^Cre^;Tau^nLacZ^*and *Phox2b^Cre^;Tau^nLacZ^* mice. In these animals neurons that expressed or express Wnt1 (in *Wnt1^Cre^;Tau^nLacZ^*) or Phox2b (in *Phox2b^Cre^;Tau^nLacZ^*), will express a nuclear β-galactosidase. **(F)** Immunohistology experiments in the jugular ganglia of *Wnt1^Cre^;Tau^nLacZ^*mice at P4. Immunohistology against β-gal (green) and Tlx3 (red, left), quantification shows that 95.7±2.6% of β-gal+ cells are Tlx3+, n=3. Immunohistology against β-gal (green) and TrkA (red, left), quantification shows that 81.8±5.0% of β-gal+ cells are TrkA+, n=3. **(G)** Immunohistology experiments in the nodose ganglia of *Phox2b^Cre^;Tau^nLacZ^* mice at P4. Immunohistology against β-gal (green) and Tlx3 (red, left), quantification shows that 99.6±0.4% of β-gal+ cells are Tlx3+, n=3. Immunohistology against β- gal (green) and TrkB (red, left), quantification shows that 99.0±1.4% of β-gal+ cells are TrkB+, n=3.

We next asked whether the Sox10+ cells with a history of Phox2b expression in the nodose ganglia belong to a lineage distinct from the Wnt1+ neural crest lineage. For this, we performed a second lineage tracing experiment using the *Wnt1^Cre^* mouse line together with the *R26^nGFP^*reporter (Figure 2A). The proportion of Phox2b+ and Sox10+ cells deriving from Wnt1+ neural crest progenitors was determined at P4. Analysis of *Wnt1^Cre^;R26^nGFP^* mice revealed that GFP was expressed in 99±0.7% (n=3) of Sox10+ glial cells in the nodose ganglion, compared to only 2.4±3.0% (n=3) of Phox2b+ neurons (Figure 2D). This result confirms previous data showing that all glial, but not neuronal, cells in the nodose ganglia are neural crest-derived, and thus the Sox10+Phox2b+ cells belong to the neural crest, and not placodal, lineage.

To assess whether our *Phox2b^Cre^* mouse line was aberrantly active in neural crest derivatives and incorrectly labeled 35.2±2.3% of Sox10+ nodose glial cells (Figure 2C middle), we compared the contributions made by the neural crest and epibranchial placodes to the neuronal populations of the jugular and nodose ganglia. To this end, we carried out long-term lineage tracing experiments using either *Wnt1^Cre^* or *Phox2b^Cre^*in combination with the reporter allele *Tau^nLacZ^* that expresses nuclear β-galactosidase (β-gal) protein exclusively in postmitotic neurons following Cre-mediated recombination (see Figure 2E for a schematic display of the genetic strategy). We first analyzed β-gal+ cells in the jugular and nodose ganglia of *Wnt1^Cre^;Tau^nLacZ^* and *Phox2b^Cre^;Tau^nLacZ^*mice and observed that the vast majority of β-gal+ cells co-expressed Tlx3 (Figure 2F left, 95.7±2.6%, n=3; Fig. 3G left, 99.6±0.4%, n=3). Thus confirming that our genetic strategy specifically labeled neuronal derivatives. In the jugular ganglion, 81.8±5.1% of β-gal+ neurons co-expressed TrkA in *Wnt1^Cre^;Tau^nLacZ^* mice (Figure 2F right, n=3), whereas virtually all nodose neurons expressed TrkB in *Phox2b^Cre^;Tau^nLacZ^* animals (Figure 2G right, 99.0±1.4%, n=3). These experiments demonstrated that, as expected, TrkA+ jugular neurons derive from the Wnt1+ neural crest, while TrkB+ nodose neurons arise from placodally-derived Phox2b+ neuroblasts. Taken together, we conclude that two distinct neural crest Wnt1+ lineages contribute to vagal ganglia derivates: i) a Wnt1+Phox2b- population that differentiates into neurons and glial cells of the jugular ganglia, as well as to roughly 60% of nodose glial cells, and ii) a Wnt1+Phox2b+ population that selectively generates roughly 40% of nodose glial cells.

**Figure 3.**
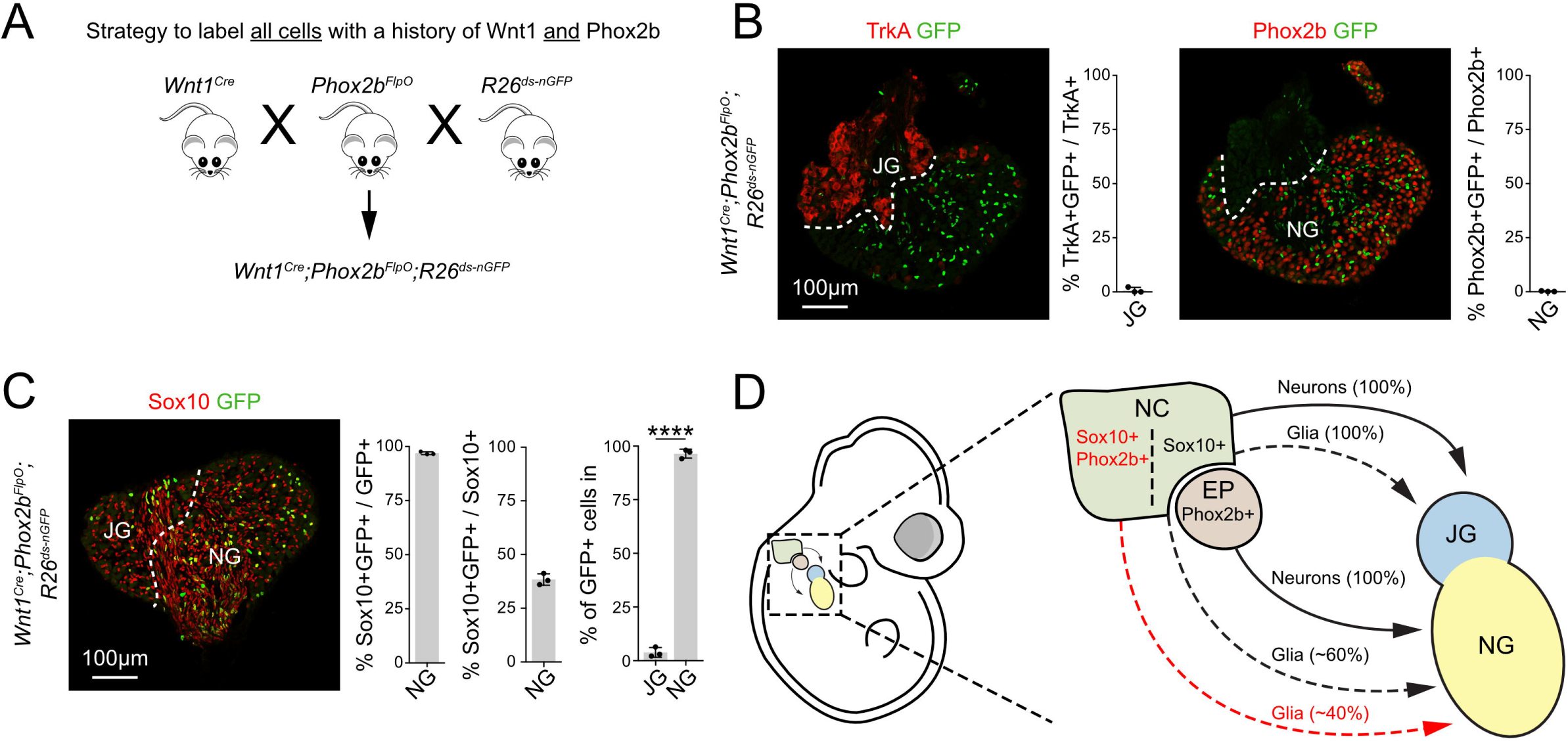
Nodose glial cells with a history of Phox2b expression are crest- derived. **(A)** To specifically label the progeny from all cells that have a history of both Wnt1 and Phox2b expression we used a reporter mouse that expresses nuclear GFP from the Rosa locus only after Cre- and Flp-mediated stop cassette excision (*R26^ds-nGFP^*) together with *Wnt1^Cre^* and *Phox2b^FlpO^*. **(B)** We examined the vagal ganglia from *Wnt1^Cre^;Phox2b^FlpO^;R26^ds-nGFP^*at P4 and performed immunohistology against GFP (green), together with TrkA (red, left) and Phox2b (red, right). Almost no TrkA+ jugular neurons (0.8±1.3%, n=3) or Phox2b+ nodose neurons (0.2±0.2%, n=3) were GFP+. **(C)** Immunohistology against GFP (green) and Sox10 (red) in *Wnt1^Cre^;Phox2b^FlpO^;R26^ds-nGFP^* at P4 revealed that 96.9±0.7% of all GFP+ cells were Sox10+ and 38.4±2.6% of all nodose Sox10+ cells were GFP+, n=3. GFP+ cells were located in the nodose, rather than the jugular ganglia (96.4±2.1% vs. 3.8±2.2%, p<0.0001, n=3). **(D)** Schema showing vagal ganglia development. The neural crest (NC, olive green) contains Sox10+ progenitors that give rise to neurons (solid lines) and glial cells (dotted lines) of the jugular ganglia (JG, blue), and the newly described Sox10+Phox2b+ progenitors that give rise to glia (dotted red line) of the nodose ganglia (NG, yellow). NG neurons derive from the Phox2b+ epibranchial placode (EP, red). Data are represented as mean ± SD, ****p < 0.0001, unpaired two-tailed t-test (J). Abbreviations: JG, jugular ganglia; NG, nodose ganglia; NC, neural crest; EP, epibranchial placode.

To better visualize the contribution of Wnt1+Phox2b+ progenitors to the nodose ganglia, we designed an intersectional genetic lineage-tracing strategy to mark all cells with a history of Wnt1 and Phox2b expression. We generated *Wnt1^Cre^*;*Phox2b^FlpO^* double transgenic animals and crossed them together with the reporter *R26^ds-nGFP^* line that expresses nuclear GFP after Cre- and Flp-mediated recombination (see Figure 3A for a schematic display of the genetic strategy). Vagal ganglia from *Wnt1^Cre^;Phox2b^FlpO^;R26^ds-nGFP^* mice were analyzed at P4 using immunohistology. We first examined whether any TrkA+ jugular or Phox2b+ nodose neurons were GFP+, and found that GFP+ cells rarely expressed either of these neuronal markers (Figure 3B, n=3). In the nodose ganglia, 96.9±0.7% of GFP+ cells co-expressed Sox10 and 38.4±2.6% of Sox10+ cells expressed GFP (Figure 3C, n=3). Furthermore, the overwhelming majority of GFP+ cells (96.4±2.1%) were located in the nodose ganglia, while the remaining GFP+ cells (3.8±2.3%) located to the TrkA+ jugular ganglia (Figure 3C, n=3). We conclude that roughly 40% of Sox10+ glial cells in the nodose ganglia derive from a neural crest progenitor population that transiently expresses Phox2b (summarized in Figure 3D).

### scRNA-seq reveals that nodose glial cells with a history of Phox2b contribute to all major glial cell subtypes in the nodose ganglia

Our data from early embryonic development indicated that a proportion of Sox10+ cells located along the vagus nerve and in the nodose anlage transiently express Phox2b between E10.5-12.5 (Figure 1). Previous work demonstrated that Sox10+Phox2b+ parasympathetic nerve-associated crest progenitors generate Schwann cells (Dyachuk et al., 2014; Espinosa-Medina et al., 2014), although it is unknown whether such progenitors can give rise to other glial cell types. To investigate which postnatal glial cell types are generated from Sox10+Phox2b+ progenitors, we used *Phox2b^Cre^;Ai14* mice to drive the expression of tdTomato fluorescent protein in all cells with a history of Phox2b expression. This strategy permanently labeled all derivatives from i) the epibranchial placode (nodose neurons displaying sustained Phox2b expression) and ii) Sox10+Phox2b+ crest-derived cells (glial cells displaying transient Phox2b expression) with cytoplasmic tdTomato protein (Figure 4A). We dissected the vagal ganglia from *Phox2b^Cre^;Ai14* mice at P4, and dissociated them into a single cell suspension using a neuronal depletion protocol (Figure 4A; see Methods). Next, we used flow cytometry to sort 384 tdTomato+ cells and generated libraries for next generation sequencing using the low-throughput, high sensitivity CEL- Seq2 method (Figure 4A) (Hashimshony et al., 2016). After sequencing 289 (75%) cells passed quality control (see methods). We detected a median of 4,484 genes and 11,900 unique molecular identifiers per cell (Figure 4B).

**Figure 4.**
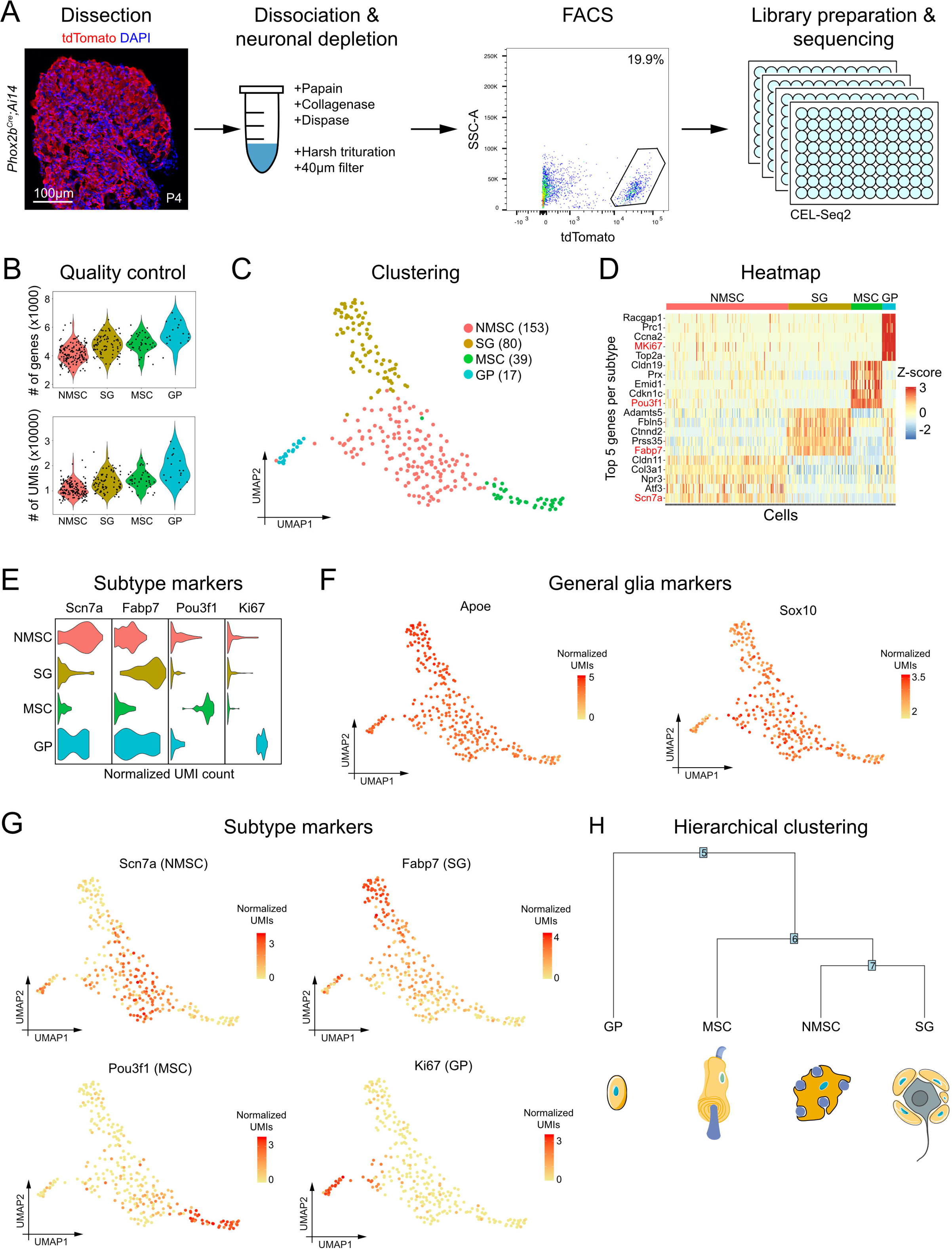
Bioinformatics analysis reveals that Sox10+Phox2b+ cells give rise to all major glial cell types in the nodose ganglia. **(A)** In order to examine the derivatives from Sox10+Phox2b+ cells we used *Phox2b^Cre^* to drive the expression of a cytoplasmic tomato protein upon Cre-mediated excision of a stop cassette. Immunofluorescence at P4 showed many tdTomato+ cells in the nodose ganglion. To investigate the non-neuronal derivatives from Sox10+Phox2b+ cells we dissociated the vagal ganglia from *Phox2b^Cre^;Ai14* mice at P4, and employed a neuronal depletion protocol. After enzymatic digestion at 37°C, ganglia were harshly mechanically triturated and passed twice through a 40µm filter. 384 tdTomato+ cells were then sorted into four 96-well plates using flow cytometry. scRNA-seq libraries were prepared using the CEL-Seq2 protocol and sequenced. **(B)** We detected a median of 4,484 genes and 11,900 unique molecular identifiers (UMIs) per cell. **(C)** UMAP plot of non-neuronal Phox2b derivatives, with each subtype labeled in a different color. The UMAP plot revealed four transcriptomically unique non-neuronal subtypes: Non-Myelinating Schwann Cells (NMSC, peach), Satellite Glia (SG, gold), Myelinating Schwann Cells (MSC, green), and Glial Precursors (GP, blue). **(D)** Heatmap showing the top 5 subtype specific genes for each subtype, ranked by p-value. Note the color code on top of the heatmap. The marker gene used to identify each subtype is shown in red. **(E)** Violin plots showing the subtype markers. *Scn7a* labels NMSC, *Fabp7* labels SG, *Pou3f1* labels MSC, and *Ki67* labels GP. **(F)** UMAP plots for general glia markers *Apoe* and *Sox10* showing that all cells in our analysis are glia. **(G)** UMAP plots for each subtype marker. **(H)** Hierarchical clustering dendrogram reveals that NMSC and SG are the most transcriptomically similar subtypes, while MSC and GP are different from each other and also from NMSC and SG. Illustrations were adapted from bioicons.com and scidraw.io and licensed under CC-BY 3.0 and CC-BY 4.0.

Further analysis distinguished four glial cell subtypes: non-myelinating Schwann cells (NMSC, 153 cells, 53% of total), satellite glia (SG, 80 cells, 28% of total), myelinating Schwann cells (MSC, 39 cells, 13% of total) and glial precursors (GP, 17 cells, 6% of total) (Figure 4C). The four glial subtypes were identified using differential expression of known marker genes (described below). Thus, the Sox10+Phox2b+ neural crest-derived glial progenitors give rise to all major glial cell subtypes in the nodose ganglia.

We identified subtype specific marker genes by performing differential gene expression analysis for each subtype (Tables 1 and 2). The top five marker genes per subtype ranked by average log fold change are displayed in a heatmap, and the genes used for further characterization are shown in red (Figure 4D). Violin plots for the subtype markers are shown in Figure 4E. All cells expressed high levels of pan-glial markers *Apoe* and *Sox10*, confirming that our approach specifically purified glial cells (Figure 4F) (Boyles et al., 1985; Kuhlbrodt et al., 1998; Britsch et al., 2001; Duan et al., 2007; Zhou et al., 2019).

Non-myelinating Schwann cells made up the largest group of cells (53% of all cells) and were defined by their expression of *Scn7a* (*Nav2.1* or *Na_x_*), *Ngfr* (*P75*) and *L1cam* (Figure 4G; Table 1) (Watanabe et al., 2002; Wolbert et al., 2020; Tasdemir-Yilmaz et al., 2021). Gene ontology (GO) term analysis revealed that the NMSC subtype was enriched for the terms “extracellular matrix organization”, “regulation of cell adhesion” and “axon ensheathment”, suggesting that they provide structural support to nodose neurons (Figure 5).

**Figure 5.**
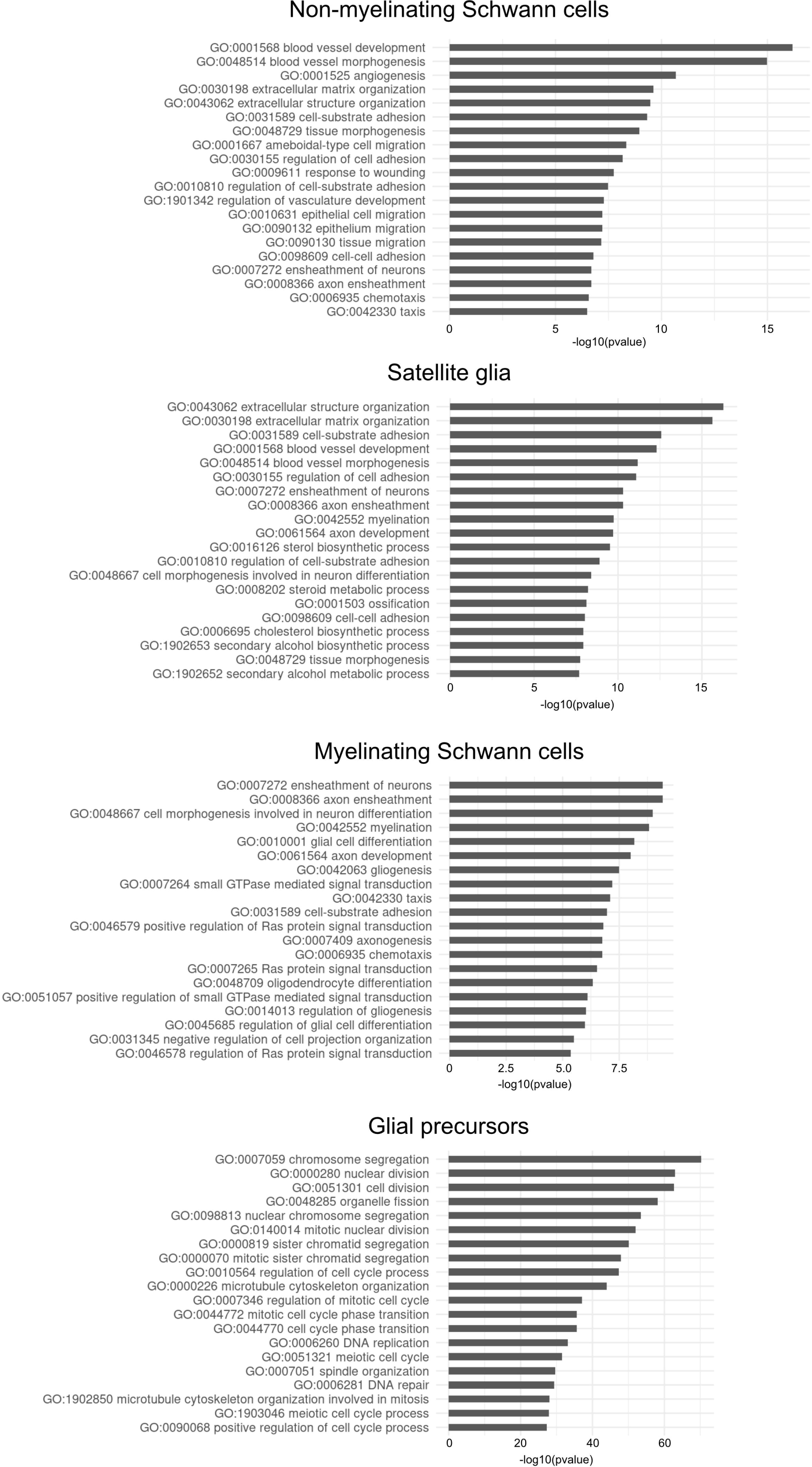
GO term analysis reveals putative roles for the four glial cell subtypes. NMSC enriched GO terms include extracellular matrix organization, cell-substrate adhesion and blood vessel development, suggesting that these cells play a role in organizing the structure of the nodose ganglia. SG are likely important for neuronal metabolism as they are enriched for sterol biosynthetic process, cholesterol biosynthetic process and secondary alcohol biosynthetic process GO terms. MSC GO terms include axon ensheathment, myelination and axonogenesis. GP are enriched for GO terms related to proliferation such as chromosome segregation, cell division and regulation of cell cycle processes.

Satellite glia, comprising 28% of the sequenced cells, expressed known satellite glia markers such as *Fabp7* (*BLBP* or *BFABP*), *Adamts5* and *Ctnnd2* (Figure 4G; Table 1) (Kurtz et al., 1994; Avraham et al., 2020, 2022; Tasdemir-Yilmaz et al., 2021). GO term analysis for enriched biological processes revealed “extracellular structure organization”, “ensheathment of neurons” and terms related to metabolism such as “sterol biosynthetic process”, “steroid metabolic process”, “cholesterol biosynthetic process” and “secondary alcohol metabolic process” (Figure 5). Thus, SG seem to be the peripheral equivalent of astrocytes, a central nervous system glial cell type that performs a homeostatic role in the brain (Hanani and Verkhratsky, 2021).

Myelinating Schwann cells, forming 13% of all cells, expressed well-characterized marker genes such as *Pou3f1* (*Oct6*), *Prx* (*Periaxin*) and *Mag* (*Myelin-associated glycoprotein*) (Figure 4G; Table 1) (Bermingham et al., 1996; Jaegle et al., 1996). GO term analysis revealed terms such as “axon ensheathment”, “ensheathment of neurons” and “myelination” (Figure 5).

The smallest subtype, glial precursors, comprised 12% of all cells and was enriched for genes typically expressed in proliferating cells such as *MKi67* (*Marker of Proliferation Ki-67*), *Top2a* (*DNA Topoisomerase II Alpha*), *Cenpe* and *Cenpf* (*Centromere Protein E and F*) (Figure 4G; Table 1). Lastly, GO term analysis confirmed that these cells are highly proliferative as the top enriched terms were “chromosome segregation”, “nuclear division” and “cell division” (Figure 5).

In summary, our bioinformatics analyses revealed that all glial cell types, including Schwann cells, satellite glia and glial precursors, derive from Sox10+Phox2b+ neural crest progenitors (Figure 4). We further identified marker genes for each subtype: NMSC (*Scn7a*), SG (*Fabp7*), MSC (*Pou3f1*), and GP (*Ki67*), and GO term analysis revealed their putative functions. Lastly, we employed hierarchical clustering to examine the relationships between glial subtypes (Figure 4H). We found that GP are the most transcriptomically unique cell type, while NMSC and SG are the most similar to one another. This reflects their biology as GP and MSC have specialized roles in proliferation and myelination respectively, while NMSCs and SG have overlapping functional roles as both provide metabolic and structural support to neurons (Harty and Monk, 2017; Hanani and Spray, 2020; Hanani and Verkhratsky, 2021).

### In vivo characterization of Phox2b derived glial cell subtypes in the nodose ganglia

Our scRNA-seq analysis (Figure 4) showed that the four subtypes of nodose glial cells derive from Sox10+Phox2b+ neural crest progenitors. To confirm this *in vivo* we performed smFISH on vagal ganglia sections from *Phox2b^Cre^;R26^nGFP^* mice at P4 (Figure 6). First, we confirmed that glial cells in the nodose ganglion have a history of Phox2b expression by combining immunohistology against GFP (green) together with smFISH to detect *Sox10* and *Phox2b* mRNA (red, Figure 6A). This revealed that many GFP+ cells with a history of Phox2b are glial cells as they express *Sox10*, but not *Phox2b*, mRNA (arrowheads in Figure 6A). Next, we used immunohistology against GFP (green) together with smFISH probes against our four glial subtype markers (red): *Scn7a* to label NMSC, *Fabp7* to label SG, *Pou3f1* to label MSC and *Ki67* to label GP (Figure 6B). Double positive cells that were GFP+Marker+ demonstrate that all four glial subtypes derive from progenitors with a history of Phox2b expression (arrowheads, Figure 6B). We additionally tested whether there was any overlap between the glial subtypes *in vivo* using dual smFISH. Simultaneous expression of two subtype markers was exceedingly rare (arrowheads, Figure 6C). Note that we could not test for overlap between *Scn7a* and *Fabp7,* as these probes were incompatible. Our bioinformatics analysis uncovered additional genes specifically expressed in each glial subtype (Figure 6D). Our data show that all four glial subtypes that derive from Sox10+Phox2b+ neural crest progenitors are present *in vivo* in nodose ganglia at P4 (Summarized in Figure 6E).

**Figure 6.**
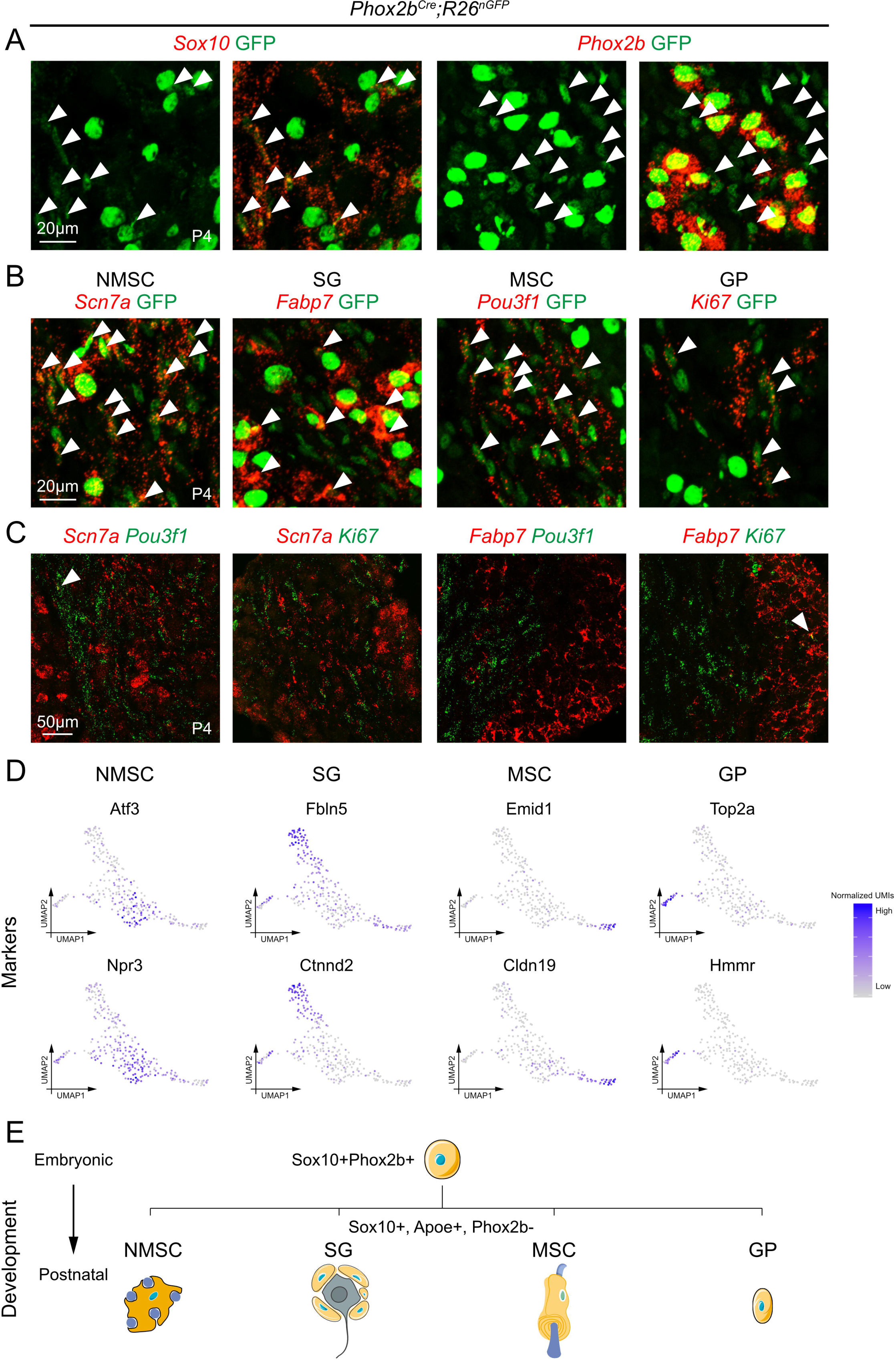
smFISH confirms that Sox10+Phox2b+ cells give rise to all major glial cell types in the nodose ganglia *in vivo*. **(A)** *Phox2b^Cre^;R26^nGFP^* mice were used to label all cells with a history of Phox2b expression with nuclear GFP (green). Nodose ganglia at P4 were analyzed using immunofluorescence against GFP (green) and smFISH probes against Sox10 (red, left) to label glial cells, and Phox2b (red, right) to label neurons. Notice the many GFP+ glial cells that are Phox2b-. Arrowheads mark GFP+ glial cells with a history of Phox2b expression that are Sox10+ (left), and Phox2b- (right). **(B)** Nodose ganglia were analyzed at P4 using immunohistology against GFP (green) together with smFISH probes (red) against *Scn7a* to label NMSC, *Fabp7* to label SG, *Pou3f1* to label MSC and *Ki67* to label GP. Arrowheads mark marker+GFP+ cells. Note that Phox2b+ nodose neurons have large, bright and round nuclei, while glial cells with a history of Phox2b expression have smaller, dimmer, oblong shaped nuclei. **(C)** smFISH with probes against glial subtype markers show that there is hardly any overlap between glial subtypes. Arrowheads label rare cells that simultaneously express subtype markers for 2 glial subtypes. **(C)** UMAPs showing additional subtype marker genes for non-myelinating Schwann cells, satellite glia, myelinating Schwann cells and glial precursors. **(D)** Summary diagram showing that Sox10+Phox2b+ cells during early development give rise to Sox10+Apoe+Phox2b- NMSC, SG, MSC and GP postnatally. Illustrations were adapted from bioicons.com and scidraw.io and licensed under CC-BY 3.0 and CC-BY 4.0.

## Discussion

The cranial neural crest generates the peripheral glial cells that populate the nodose ganglia and support resident viscerosensory neurons in their homeostatic functions. Here, we show that some Sox10+ neural crest-derived precursors transiently express Phox2b, before populating the nodose, but not the jugular ganglia, with glial cells. Glial derivatives from the Sox10+Phox2b+ neural crest progenitors include myelinating and non-myelinating Schwann cells, as well as satellite glia and glial precursors. scRNA-seq revealed the transcriptomic signatures of these four major glial cell types, which we confirm derive from progenitors with a history of Phox2b expression. Our work demonstrates the existence of a novel cranial neural crest progenitor population, namely nerve-associated Sox10+Phox2b+ cells, that generate the entire repertoire of nodose, but not jugular, glial subtypes.

### Nodose glial cell development

From the discovery of the neural crest by His in 1868, this progenitor niche has captured the attention of generations of scientists given the vast multitude of cell types that it generates (His, W., 1868; Dupont, 2018; Tang and Bronner, 2020). The cranial neural crest gives rise to many different cells that populate the head and face region in vertebrates, including bone, cartilage, melanocytes, as well as sensory neurons and glial cells (Le Douarin et al., 2004; Erickson et al., 2023). Until the development of quail-chick transplantation as a means of tracing the derivatives from different embryonic progenitor zones, there was much debate over the exact development of cranial sensory ganglia (Le Douarin, 1986). The generation of nodose ganglia was particularly controversial and it was only in 1980 that its glial component was definitively shown to derive from the neural crest, and its neurons from the epibranchial placode (Narayanan and Narayanan, 1980).

Although the same cranial neural crest gives rise to glial cells in both the nodose and the neighboring jugular ganglia, it is only in the nodose ganglia where some glial cells have a history of transient Phox2b expression (Figure 3). Transient Phox2b expression in Sox10+ cells has been described in other crest-derived progenitors that choose between neuronal or glial fates such as in enteric and parasympathetic ganglia (Dyachuk et al., 2014; Espinosa-Medina et al., 2014, 2017; Soldatov et al., 2019). In contrast to these other autonomic structures that derive their neuronal and glial components from the neural crest, neural crest only contributes glial cells to the nodose ganglia. Recent scRNA-seq studies revealed that in the neural crest, cell fate is decided by a series of binary decisions between competing transcriptional programs (Soldatov et al., 2019; Faure et al., 2020). Thus, Phox2b is expressed early in autonomic crest progenitors that differentiate into neurons and glial cells, but its expression is quickly downregulated in glial cells and only maintained in neurons (Dyachuk et al., 2014; Espinosa-Medina et al., 2014). Why Phox2b, a master regulator of autonomic cell fate, is expressed in the cranial neural crest progenitors that generate Phox2b- glial cells in the nodose ganglia was not immediately apparent. It is now known that sympathetic and enteric, but not somatosensory, neurons depend on Phox2b for their development (Pattyn et al., 1999; Espinosa-Medina et al., 2017; Soldatov et al., 2019). Further, nerve associated crest progenitors along the vagus nerve generate Phox2b+ neurons that reside in esophageal and stomach enteric ganglia (Espinosa-Medina et al., 2017). Lastly, genetic studies showed that Phox2b suppresses somatosensory fates in favor of viscerosensory ones (D’Autréaux et al., 2011). Thus, it seems as though the crest-derived Sox10+Phox2b+ progenitors that give rise to nodose glial cells described here are part and parcel of the same population of crest-derived cells that have the potential to generate other Phox2b+ autonomic neurons, but not Phox2b- somatosensory neurons. We captured the Sox10+Phox2b+ cells at a cross- roads, deciding between turning off Phox2b expression to generate the full complement of nodose glial cells, or maintaining Phox2b expression and continuing to travel along the vagus nerve to generate sympathetic or enteric neurons.

One of the major questions regarding the multipotent neural crest is whether cell fates are determined intrinsically based on their lineage, or if extrinsic factors also play a role (Le Douarin, 1986). A quail-chick transplantation study by Ayer-le Lievre and Le Douarin addressed this question by grafting quail crest into a chick host, letting the host develop, and then removed the chimeric nodose ganglia and back-transplanted it into the cranial neural crest of a younger chick host (Ayer-Le Lievre and Le Douarin, 1982). When they examined the chick hosts that received chimeric quail-chick nodose ganglia grafts, they found quail cells in numerous sympathetic and enteric, but not somatosensory, ganglia (Ayer-Le Lievre and Le Douarin, 1982). Thus, the cranial neural crest that *in vivo* only generates nodose glial cells, contains numerous “unused” autonomic neural potentialities, demonstrating that extrinsic factors play a role in neural crest fate selection.

The transient expression of Phox2b in some Sox10+ crest-derived progenitors that generate nodose, but not jugular, glial cells might be due to environmental differences. For instance, transient expression of Phox2b in nodose glial cells could be induced by interactions with other Phox2b+ cells such as viscerosensory or visceromotor neurons that form the vagus nerve, or placodal neuronal progenitors as they travel to the nodose anlage. Although the mechanism governing Phox2b expression in crest-derived nodose glial cells is unclear, the “unused” fates of the crest cells described by Ayer-le Lievre and Le Douarin that become available in a new environment are only those that depend on the expression of Phox2b.

### Glial cell diversity

The development of scRNA-seq has uncovered the remarkable heterogeneity of cell-types contained within the central and peripheral nervous systems. More recently, this technique has revealed glial heterogeneity in peripheral nerves such as the brachial plexus and sciatic nerve, as well as in peripheral ganglia such as the dorsal root ganglia, spiral ganglia, and superior cervical ganglia (Wolbert et al., 2020; Tasdemir-Yilmaz et al., 2021; Avraham et al., 2022; Mapps et al., 2022). While the glial cell types found in peripheral nerves only consist of myelinating and non-myelinating Schwann cells, peripheral ganglia also include satellite glial cells (Jessen and Mirsky, 2005). These different types of peripheral glial cells work to provide trophic support and myelination to neurons in the peripheral nervous system (Jessen and Mirsky, 2005; Jessen et al., 2015; Hanani and Spray, 2020). Here, we show that neural-crest-derived Sox10+Phox2b+ progenitors give rise to satellite glia and glial precursors, in addition to Schwann cells.

While recent studies profiled the transcriptomes of glial cells found in sympathetic, auditory and somatosensory ganglia (Tasdemir-Yilmaz et al., 2021; Mapps et al., 2022), here we sequenced glial cells from the viscerosensory nodose ganglia. Interestingly, although the neurons of these different ganglia carry out vastly different functions, they are supported by the same four cardinal glial cell types: non-myelinating and myelinating Schwann cells, satellite glia, and glial precursors. Thus, peripheral glial cells converge on the same core cell types to support different neurons that mediate homeostatic functions (nodose viscerosensory neurons), hearing (spiral ganglia neurons) and somatosensation (dorsal root ganglia neurons).

## Acknowledgements

E.D.L. was supported by an MDC Internal PhD Fellowship. This work was also supported by the Deutsche Forschungsgemeinschaft (DFG, German Research Foundation) under Germanýs Excellence Strategy – EXC- 2049 – 39068808 and by the Helmholtz Association. We thank Petra Stallerow and Claudia Päseler for animal husbandry. The authors thank Carmen Birchmeier (MDC, Berlin), Fritz Rathjen (MDC, Berlin), Joscha Griger (TUM, Munich) and Jean-François Brunet (École normale supérieure, Paris) for critically reading the manuscript. The authors thank Hans-Peter Rahn of the MDC Flow Cytometry facility and the MDC Next Generation Sequencing facility.

## Author contributions

E.D.L. designed research, performed research, analyzed data, and wrote the paper. A.M. performed research and analyzed data. S.B. and P.L.R. performed research.

---

**Extended Data Figure 1-1. Sox10+ cells in the nodose anlage are non-neuronal.** Immunohistology against Sox10 (green) and Tlx3 (red) in the nodose ganglion at E12.5. Boxed region is shown magnified below and a Sox10+ nucleus is circled with a dotted line. Quantifications on the right show that 0.1±0.1% and 0.0±0.0% of Sox10+ cells co-express Tlx3 at E11.5 and E12.5, respectively.

