## Supplementary figures and images for "Molecular characterization of nodose ganglia development reveals a novel population of Phox2b+ glial progenitors in mice"

### Extended Data Figure 1-1

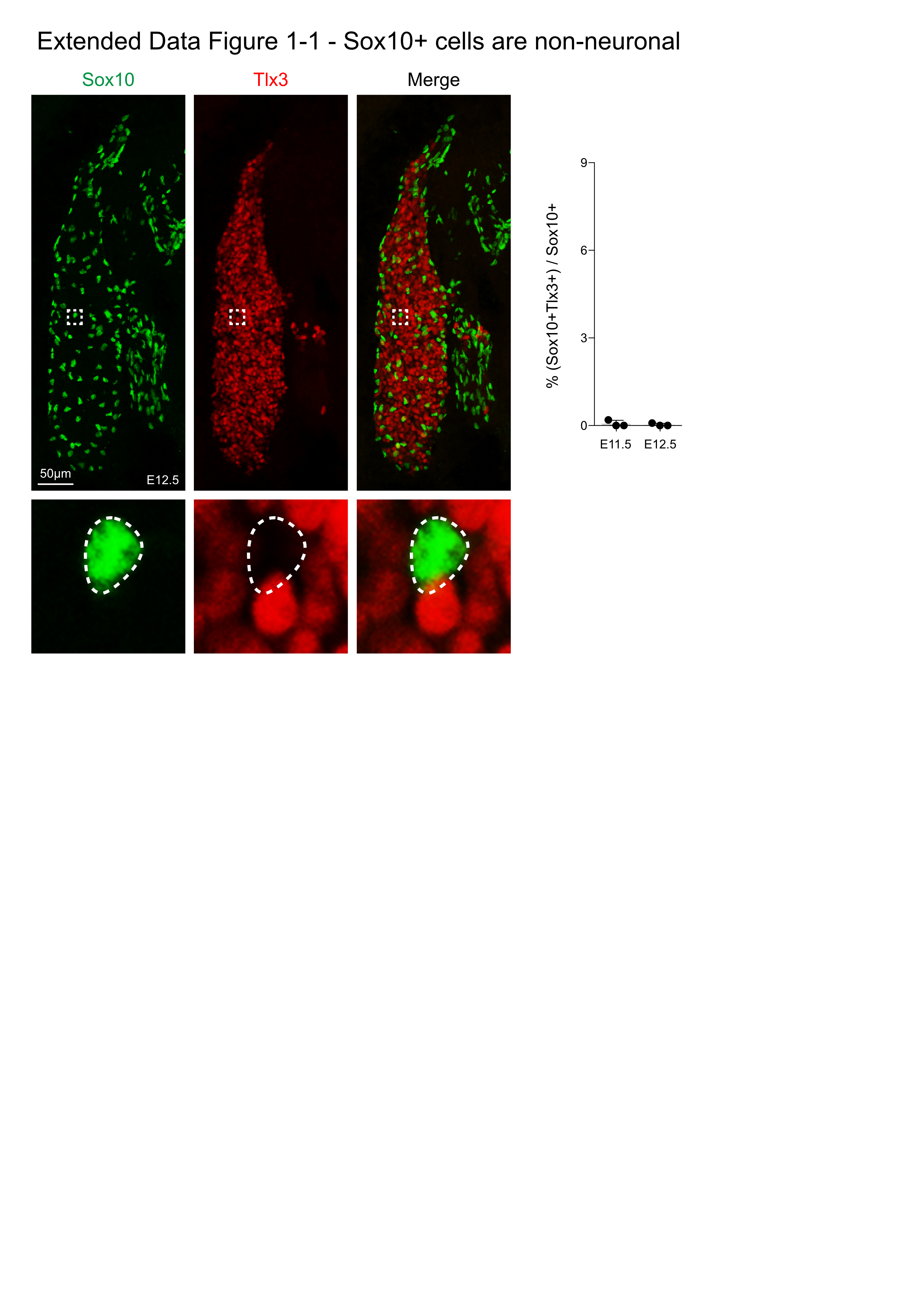
