## Supplementary material for "Molecular characterization of nodose ganglia development reveals a novel population of Phox2b+ glial progenitors in mice": Table 1

**Supplementary Table 1. Top 25 positive marker genes for each cluster ranked by average log fold change.**

| **Gene name** | **Average log fold change** | **P value** | **P value adjusted** | **pct.1** | **pct.2** | **Cluster** |
| --- | --- | --- | --- | --- | --- | --- |
| Scn7a | 1.3454415 | 7.73E-23 | 1.14E-18 | 0.876 | 0.5 | NMSC |
| Atf3 | 1.1848592 | 1.30E-10 | 1.93E-06 | 0.667 | 0.434 | NMSC |
| Npr3 | 1.0604613 | 6.34E-19 | 9.39E-15 | 0.882 | 0.669 | NMSC |
| Col3a1 | 0.8784661 | 2.77E-25 | 4.10E-21 | 1 | 1 | NMSC |
| Cldn11 | 0.8486067 | 5.81E-19 | 8.59E-15 | 0.869 | 0.5 | NMSC |
| Zfp36 | 0.8384565 | 2.01E-13 | 2.97E-09 | 0.771 | 0.463 | NMSC |
| Ngfr | 0.8317061 | 4.64E-14 | 6.87E-10 | 0.85 | 0.618 | NMSC |
| Jun | 0.8278197 | 7.68E-22 | 1.14E-17 | 1 | 0.985 | NMSC |
| Wwp1 | 0.8235066 | 7.37E-22 | 1.09E-17 | 0.954 | 0.794 | NMSC |
| Junb | 0.8211079 | 2.38E-16 | 3.52E-12 | 0.941 | 0.765 | NMSC |
| Socs3 | 0.7972627 | 8.49E-13 | 1.26E-08 | 0.948 | 0.713 | NMSC |
| Btg2 | 0.7859394 | 2.25E-12 | 3.34E-08 | 0.758 | 0.441 | NMSC |
| Fosb | 0.774052 | 2.42E-14 | 3.59E-10 | 0.915 | 0.735 | NMSC |
| Klf6 | 0.7573566 | 3.07E-14 | 4.54E-10 | 0.98 | 0.949 | NMSC |
| Nrp2 | 0.7572285 | 4.28E-22 | 6.34E-18 | 0.974 | 0.912 | NMSC |
| Egr1 | 0.7443666 | 4.73E-26 | 7.00E-22 | 0.993 | 0.963 | NMSC |
| Ier2 | 0.7188142 | 6.02E-13 | 8.91E-09 | 0.922 | 0.721 | NMSC |
| Fos | 0.7114711 | 3.27E-24 | 4.84E-20 | 1 | 0.993 | NMSC |
| Sbspon | 0.706978 | 2.75E-17 | 4.08E-13 | 0.869 | 0.522 | NMSC |
| Nr4a1 | 0.6605907 | 8.38E-15 | 1.24E-10 | 1 | 0.956 | NMSC |
| Abca8b | 0.6576159 | 3.27E-21 | 4.85E-17 | 0.993 | 0.912 | NMSC |
| Col1a1 | 0.6446967 | 3.17E-11 | 4.70E-07 | 0.98 | 0.904 | NMSC |
| Itgb4 | 0.6420016 | 5.93E-18 | 8.78E-14 | 0.948 | 0.809 | NMSC |
| L1cam | 0.6322598 | 6.41E-16 | 9.49E-12 | 0.98 | 0.772 | NMSC |
| Mmp2 | 0.6232632 | 4.23E-10 | 6.26E-06 | 0.51 | 0.221 | NMSC |
| Fabp7 | 1.8190219 | 1.14911E-27 | 1.70E-23 | 1 | 0.746 | SG |
| Prss35 | 1.4234951 | 4.46843E-32 | 6.62E-28 | 1 | 0.703 | SG |
| Ctnnd2 | 1.3588487 | 6.71187E-29 | 9.94E-25 | 0.912 | 0.321 | SG |
| Fbln5 | 1.2223476 | 7.35366E-22 | 1.09E-17 | 0.95 | 0.646 | SG |
| Adamts5 | 1.1884403 | 4.0509E-23 | 6.00E-19 | 1 | 0.876 | SG |
| Epas1 | 1.1763771 | 1.48374E-20 | 2.20E-16 | 0.7 | 0.182 | SG |
| Hey2 | 1.0953341 | 8.4934E-24 | 1.26E-19 | 0.8 | 0.278 | SG |
| Sfrp5 | 1.0753977 | 3.49345E-23 | 5.17E-19 | 0.988 | 0.847 | SG |
| Igfbp4 | 1.0093095 | 9.32092E-23 | 1.38E-18 | 0.975 | 0.656 | SG |
| Fbln2 | 0.9628845 | 5.52588E-22 | 8.18E-18 | 1 | 0.981 | SG |
| Fgfr1 | 0.9430148 | 3.91708E-21 | 5.80E-17 | 0.962 | 0.871 | SG |
| Ptprz1 | 0.9213493 | 1.75032E-24 | 2.59E-20 | 1 | 0.866 | SG |
| Fmo1 | 0.9144529 | 1.33537E-26 | 1.98E-22 | 0.762 | 0.144 | SG |
| Cdh11 | 0.9121881 | 4.73561E-23 | 7.01E-19 | 0.988 | 0.871 | SG |
| Hmgcs1 | 0.9117534 | 1.60181E-13 | 2.37E-09 | 0.975 | 0.856 | SG |
| Mest | 0.910458 | 9.52418E-20 | 1.41E-15 | 0.85 | 0.364 | SG |
| Mapre2 | 0.9067151 | 3.4386E-24 | 5.09E-20 | 1 | 0.947 | SG |
| Me1 | 0.9043683 | 8.02962E-24 | 1.19E-19 | 0.962 | 0.603 | SG |
| Kit | 0.880006 | 2.54453E-18 | 3.77E-14 | 0.838 | 0.407 | SG |
| Acsbg1 | 0.8487004 | 2.16553E-19 | 3.21E-15 | 0.95 | 0.632 | SG |
| Cst3 | 0.8456367 | 3.85576E-17 | 5.71E-13 | 1 | 1 | SG |
| Lect1 | 0.8257245 | 3.41811E-16 | 5.06E-12 | 0.55 | 0.115 | SG |
| Slc1a4 | 0.8143027 | 1.0529E-18 | 1.56E-14 | 0.725 | 0.22 | SG |
| Apoe | 0.8089571 | 4.8781E-31 | 7.22E-27 | 1 | 0.995 | SG |
| Col26a1 | 0.8028451 | 5.83969E-21 | 8.65E-17 | 0.975 | 0.598 | SG |
| Pou3f1 | 2.533831 | 2.31E-28 | 3.43E-24 | 1 | 0.332 | MSC |
| Cdkn1c | 2.369622 | 2.80E-20 | 4.14E-16 | 0.974 | 0.548 | MSC |
| Emid1 | 1.875401 | 6.17E-17 | 9.13E-13 | 0.667 | 0.148 | MSC |
| Prx | 1.810557 | 3.38E-15 | 5.00E-11 | 0.974 | 0.608 | MSC |
| Cldn19 | 1.76779 | 6.05E-24 | 8.95E-20 | 0.923 | 0.26 | MSC |
| Csrp2 | 1.52758 | 9.52E-26 | 1.41E-21 | 0.897 | 0.18 | MSC |
| Slc36a2 | 1.519632 | 1.19E-20 | 1.76E-16 | 0.462 | 0.024 | MSC |
| Pllp | 1.488328 | 2.51E-20 | 3.71E-16 | 0.897 | 0.3 | MSC |
| Kcna1 | 1.480256 | 7.30E-20 | 1.08E-15 | 1 | 0.648 | MSC |
| Ncmap | 1.459536 | 1.09E-18 | 1.62E-14 | 0.538 | 0.056 | MSC |
| Bzw2 | 1.454874 | 1.15E-19 | 1.71E-15 | 0.974 | 0.66 | MSC |
| Fxyd6 | 1.379751 | 1.81E-04 | 1.00E+00 | 0.692 | 0.548 | MSC |
| Dusp15 | 1.331891 | 1.15E-25 | 1.70E-21 | 0.744 | 0.092 | MSC |
| Peli2 | 1.281033 | 1.90E-18 | 2.82E-14 | 0.974 | 0.548 | MSC |
| Nkd1 | 1.258765 | 2.23E-20 | 3.30E-16 | 0.897 | 0.288 | MSC |
| Mag | 1.237397 | 1.38E-07 | 2.04E-03 | 0.744 | 0.396 | MSC |
| Gldn | 1.202283 | 1.68E-25 | 2.49E-21 | 0.487 | 0.012 | MSC |
| Gypc | 1.176134 | 2.65E-26 | 3.92E-22 | 0.872 | 0.16 | MSC |
| Cuedc2 | 1.169694 | 2.01E-11 | 2.97E-07 | 0.974 | 0.908 | MSC |
| Fa2h | 1.135748 | 1.18E-18 | 1.74E-14 | 0.872 | 0.252 | MSC |
| Mpp6 | 1.134361 | 3.36E-14 | 4.98E-10 | 0.744 | 0.256 | MSC |
| Nav1 | 1.128743 | 5.10E-16 | 7.54E-12 | 1 | 0.584 | MSC |
| Rasal2 | 1.120441 | 1.22E-14 | 1.81E-10 | 0.923 | 0.596 | MSC |
| Smtn | 1.081293 | 6.75E-16 | 1.00E-11 | 0.974 | 0.572 | MSC |
| Fyn | 1.078509 | 2.92E-17 | 4.33E-13 | 1 | 0.804 | MSC |
| Top2a | 2.855656 | 1.48E-17 | 2.19E-13 | 1 | 0.25 | GP |
| Mki67 | 2.570845 | 5.85E-21 | 8.67E-17 | 1 | 0.18 | GP |
| Ccna2 | 2.3057 | 2.47E-31 | 3.66E-27 | 1 | 0.081 | GP |
| Prc1 | 2.132067 | 1.88E-22 | 2.78E-18 | 1 | 0.151 | GP |
| Racgap1 | 2.055056 | 7.73E-23 | 1.14E-18 | 1 | 0.151 | GP |
| Cenpf | 2.004354 | 1.73E-18 | 2.57E-14 | 1 | 0.191 | GP |
| Ube2c | 1.969373 | 7.28E-33 | 1.08E-28 | 0.941 | 0.055 | GP |
| Smc4 | 1.943801 | 7.28E-13 | 1.08E-08 | 1 | 0.555 | GP |
| Ckap2l | 1.902747 | 6.24E-39 | 9.24E-35 | 1 | 0.048 | GP |
| Hmmr | 1.896509 | 1.38E-42 | 2.04E-38 | 1 | 0.037 | GP |
| Cdca3 | 1.860298 | 2.13E-26 | 3.15E-22 | 1 | 0.11 | GP |
| Tpx2 | 1.813751 | 5.10E-24 | 7.55E-20 | 1 | 0.136 | GP |
| Cdk1 | 1.804463 | 1.45E-24 | 2.14E-20 | 0.941 | 0.107 | GP |
| Ccnb1 | 1.80345 | 2.52E-35 | 3.74E-31 | 1 | 0.059 | GP |
| Ncapd2 | 1.759535 | 2.20E-18 | 3.26E-14 | 1 | 0.232 | GP |
| Ckap2 | 1.757167 | 5.25E-23 | 7.78E-19 | 1 | 0.14 | GP |
| H2afx | 1.711216 | 2.05E-21 | 3.04E-17 | 1 | 0.169 | GP |
| Fam64a | 1.702165 | 5.37E-41 | 7.95E-37 | 1 | 0.04 | GP |
| Cenpe | 1.693033 | 1.19E-22 | 1.76E-18 | 0.941 | 0.107 | GP |
| Aspm | 1.691371 | 8.86E-29 | 1.31E-24 | 1 | 0.088 | GP |
| Cenpa | 1.690392 | 8.88E-14 | 1.31E-09 | 0.941 | 0.265 | GP |
| Mis18bp1 | 1.689529 | 1.40E-27 | 2.07E-23 | 1 | 0.103 | GP |
| Casc5 | 1.666056 | 2.99E-39 | 4.42E-35 | 1 | 0.044 | GP |
| Pbk | 1.64447 | 1.38E-42 | 2.04E-38 | 1 | 0.037 | GP |
| Birc5 | 1.642404 | 5.72E-30 | 8.47E-26 | 1 | 0.088 | GP |
