## Supplementary material for "Molecular characterization of nodose ganglia development reveals a novel population of Phox2b+ glial progenitors in mice": Table 2

**Supplementary Table 2. Top 25 negative marker genes for each cluster ranked by average log fold change.**

| **Gene name** | **Average log fold change** | **P value** | **P value adjusted** | **pct.1** | **pct.2** | **Cluster** |
| --- | --- | --- | --- | --- | --- | --- |
| Fabp7 | -1.4920448 | 8.11E-08 | 1.20E-03 | 0.81 | 0.824 | NMSC |
| Fbln5 | -1.1724742 | 9.82E-22 | 1.45E-17 | 0.556 | 0.926 | NMSC |
| Ctnnd2 | -1.1525979 | 3.00E-14 | 4.43E-10 | 0.32 | 0.669 | NMSC |
| Prss35 | -1.0890231 | 6.57E-13 | 9.73E-09 | 0.732 | 0.846 | NMSC |
| Emid1 | -0.9415067 | 3.84E-04 | 1.00E+00 | 0.144 | 0.301 | NMSC |
| Maf | -0.9023147 | 1.14E-18 | 1.68E-14 | 0.784 | 0.978 | NMSC |
| Adamts5 | -0.8750677 | 2.62E-07 | 3.88E-03 | 0.902 | 0.919 | NMSC |
| Epas1 | -0.8502615 | 2.95E-11 | 4.37E-07 | 0.163 | 0.507 | NMSC |
| Hey2 | -0.8251426 | 2.71E-13 | 4.02E-09 | 0.248 | 0.618 | NMSC |
| Mest | -0.7657799 | 1.12E-11 | 1.66E-07 | 0.34 | 0.676 | NMSC |
| Cdh11 | -0.7449968 | 6.21E-14 | 9.20E-10 | 0.85 | 0.963 | NMSC |
| Serpine2 | -0.7440726 | 3.22E-23 | 4.77E-19 | 0.961 | 1 | NMSC |
| Sfrp5 | -0.7209466 | 1.68E-07 | 2.48E-03 | 0.856 | 0.919 | NMSC |
| Me1 | -0.7186752 | 2.77E-13 | 4.10E-09 | 0.595 | 0.824 | NMSC |
| Csrp2 | -0.7181241 | 1.19E-06 | 1.75E-02 | 0.163 | 0.404 | NMSC |
| Slc36a2 | -0.6979428 | 1.77E-04 | 1.00E+00 | 0.026 | 0.147 | NMSC |
| Tyrp1 | -0.6745873 | 2.25E-04 | 1.00E+00 | 0.137 | 0.309 | NMSC |
| Fmo1 | -0.6709603 | 6.43E-12 | 9.52E-08 | 0.15 | 0.5 | NMSC |
| Kit | -0.6708608 | 3.89E-11 | 5.76E-07 | 0.366 | 0.706 | NMSC |
| Fbln2 | -0.6659355 | 3.36E-09 | 4.98E-05 | 0.98 | 0.993 | NMSC |
| Ncmap | -0.6550502 | 1.31E-03 | 1.00E+00 | 0.065 | 0.184 | NMSC |
| Sncg | -0.6482114 | 6.85E-11 | 1.01E-06 | 0.255 | 0.61 | NMSC |
| Glul | -0.644325 | 1.96E-10 | 2.90E-06 | 0.523 | 0.787 | NMSC |
| Bzw2 | -0.6364356 | 5.96E-05 | 8.82E-01 | 0.621 | 0.794 | NMSC |
| Prc1 | -0.63628 | 6.93E-03 | 1.00E+00 | 0.144 | 0.265 | NMSC |
| Pou3f1 | -1.6159925 | 9.12E-09 | 1.35E-04 | 0.188 | 0.512 | SG |
| Cdkn1c | -1.365596 | 1.42E-06 | 2.11E-02 | 0.462 | 0.66 | SG |
| Col1a1 | -1.1920536 | 1.08E-16 | 1.60E-12 | 0.875 | 0.971 | SG |
| Fosb | -1.179236 | 8.49E-17 | 1.26E-12 | 0.688 | 0.885 | SG |
| Btg2 | -1.1252537 | 8.84E-13 | 1.31E-08 | 0.325 | 0.718 | SG |
| Prx | -1.1137562 | 2.91E-10 | 4.31E-06 | 0.412 | 0.751 | SG |
| Atf3 | -1.1050339 | 2.32E-07 | 3.43E-03 | 0.375 | 0.627 | SG |
| Scn7a | -1.0923288 | 4.73E-10 | 7.00E-06 | 0.525 | 0.766 | SG |
| Npr3 | -1.0727683 | 4.64E-12 | 6.87E-08 | 0.662 | 0.828 | SG |
| Nav1 | -1.0288453 | 6.52E-16 | 9.66E-12 | 0.35 | 0.751 | SG |
| Junb | -1.0167097 | 1.42E-15 | 2.11E-11 | 0.725 | 0.909 | SG |
| Kcna1 | -0.9827515 | 1.72E-11 | 2.54E-07 | 0.462 | 0.785 | SG |
| Col3a1 | -0.9726432 | 2.09E-18 | 3.10E-14 | 1 | 1 | SG |
| Nr4a1 | -0.9377319 | 3.63E-17 | 5.37E-13 | 0.95 | 0.99 | SG |
| Zfp36 | -0.9017113 | 2.19E-10 | 3.25E-06 | 0.412 | 0.708 | SG |
| Egr2 | -0.8818761 | 7.69E-10 | 1.14E-05 | 0.362 | 0.636 | SG |
| Dusp6 | -0.869753 | 4.11E-14 | 6.09E-10 | 0.812 | 0.909 | SG |
| Reln | -0.8638117 | 6.60E-13 | 9.77E-09 | 0.562 | 0.842 | SG |
| Socs3 | -0.8497213 | 5.21E-09 | 7.71E-05 | 0.738 | 0.876 | SG |
| Smtn | -0.842174 | 3.89E-12 | 5.76E-08 | 0.4 | 0.713 | SG |
| Fos | -0.8190388 | 2.06E-19 | 3.04E-15 | 1 | 0.995 | SG |
| Cldn19 | -0.8184983 | 4.24E-05 | 6.27E-01 | 0.2 | 0.407 | SG |
| Ier2 | -0.795103 | 5.08E-11 | 7.53E-07 | 0.65 | 0.895 | SG |
| Gjc3 | -0.7886277 | 2.00E-21 | 2.96E-17 | 0.988 | 0.995 | SG |
| Klf6 | -0.7857608 | 9.22E-09 | 1.36E-04 | 0.938 | 0.976 | SG |
| Fabp7 | -1.9355493 | 2.75E-12 | 4.07E-08 | 0.41 | 0.88 | MSC |
| Scn7a | -1.7541323 | 1.12E-09 | 1.65E-05 | 0.359 | 0.752 | MSC |
| Entpd2 | -1.6081918 | 9.10E-21 | 1.35E-16 | 0.692 | 1 | MSC |
| Ptprz1 | -1.3085331 | 1.08E-14 | 1.59E-10 | 0.538 | 0.96 | MSC |
| Abca8b | -1.3046711 | 3.80E-18 | 5.62E-14 | 0.718 | 0.992 | MSC |
| Cyr61 | -1.2275561 | 1.78E-06 | 2.63E-02 | 0.513 | 0.764 | MSC |
| Matn2 | -1.2223053 | 1.53E-17 | 2.27E-13 | 0.821 | 0.996 | MSC |
| Adamts5 | -1.2078588 | 4.26E-11 | 6.31E-07 | 0.718 | 0.94 | MSC |
| Mmd2 | -1.1816661 | 6.48E-17 | 9.59E-13 | 0.641 | 0.976 | MSC |
| Ptn | -1.1775485 | 5.37E-16 | 7.94E-12 | 0.615 | 0.976 | MSC |
| Mapre2 | -1.1618803 | 1.07E-13 | 1.58E-09 | 0.923 | 0.968 | MSC |
| Prss35 | -1.1440868 | 4.89E-08 | 7.24E-04 | 0.513 | 0.828 | MSC |
| Abca8a | -1.1424352 | 8.66E-20 | 1.28E-15 | 0.974 | 1 | MSC |
| Id3 | -1.133172 | 4.60E-11 | 6.81E-07 | 0.154 | 0.704 | MSC |
| Ctnnd2 | -1.0771028 | 1.21E-06 | 1.80E-02 | 0.154 | 0.536 | MSC |
| Slc35f1 | -1.0523023 | 7.60E-15 | 1.12E-10 | 0.769 | 0.984 | MSC |
| Cst3 | -1.0384663 | 6.91E-14 | 1.02E-09 | 1 | 1 | MSC |
| L1cam | -0.9942918 | 8.91E-11 | 1.32E-06 | 0.615 | 0.924 | MSC |
| Apoe | -0.9904277 | 9.97E-16 | 1.48E-11 | 0.974 | 1 | MSC |
| Hmgcs1 | -0.9824656 | 1.51E-08 | 2.24E-04 | 0.795 | 0.904 | MSC |
| Atp1b2 | -0.9522054 | 1.74E-11 | 2.58E-07 | 0.359 | 0.832 | MSC |
| Sfrp5 | -0.950984 | 1.01E-08 | 1.50E-04 | 0.744 | 0.908 | MSC |
| Sparcl1 | -0.9473638 | 9.02E-14 | 1.34E-09 | 0.949 | 1 | MSC |
| Ndrg2 | -0.9454483 | 5.06E-11 | 7.49E-07 | 0.641 | 0.896 | MSC |
| Cldn11 | -0.9337366 | 7.66E-08 | 1.13E-03 | 0.359 | 0.748 | MSC |
| Cdkn1c | -1.3730113 | 5.51E-03 | 1.00E+00 | 0.353 | 0.621 | GP |
| Prx | -1.1132678 | 5.75E-03 | 1.00E+00 | 0.529 | 0.665 | GP |
| Kcna1 | -1.0276728 | 3.12E-03 | 1.00E+00 | 0.471 | 0.71 | GP |
| Mxd4 | -0.9631035 | 2.28E-08 | 3.37E-04 | 0.647 | 0.915 | GP |
| Fam129a | -0.8840875 | 9.21E-04 | 1.00E+00 | 0.353 | 0.629 | GP |
| Cd59a | -0.7983102 | 2.70E-05 | 3.99E-01 | 0.824 | 0.938 | GP |
| Ypel3 | -0.7788885 | 2.37E-06 | 3.51E-02 | 0.647 | 0.912 | GP |
| Pik3ip1 | -0.7511425 | 6.43E-05 | 9.52E-01 | 0.176 | 0.64 | GP |
| Col27a1 | -0.7371896 | 3.27E-04 | 1.00E+00 | 0.824 | 0.882 | GP |
| Mpz | -0.7313429 | 1.67E-04 | 1.00E+00 | 1 | 1 | GP |
| Itih5 | -0.7218123 | 6.75E-03 | 1.00E+00 | 1 | 0.956 | GP |
| Hmgcs2 | -0.7170613 | 1.23E-06 | 1.82E-02 | 1 | 0.967 | GP |
| Plp1 | -0.7040201 | 3.05E-08 | 4.51E-04 | 1 | 1 | GP |
| Fosb | -0.6991397 | 4.37E-03 | 1.00E+00 | 0.647 | 0.842 | GP |
| Prkcd | -0.6900387 | 1.24E-04 | 1.00E+00 | 0.471 | 0.776 | GP |
| Jun | -0.6882787 | 1.10E-03 | 1.00E+00 | 0.941 | 0.996 | GP |
| Kcna2 | -0.6737272 | 8.11E-05 | 1.00E+00 | 0.941 | 0.993 | GP |
| Abca1 | -0.6708194 | 6.18E-04 | 1.00E+00 | 0.647 | 0.79 | GP |
| Egr1 | -0.6697287 | 5.41E-04 | 1.00E+00 | 0.882 | 0.985 | GP |
| Cnp | -0.6657583 | 1.25E-06 | 1.85E-02 | 1 | 1 | GP |
| Zfp651 | -0.6568236 | 2.81E-05 | 4.17E-01 | 0.412 | 0.779 | GP |
| Ccnd1 | -0.6505622 | 1.95E-03 | 1.00E+00 | 0.941 | 0.908 | GP |
| Kdm5b | -0.6462374 | 7.76E-05 | 1.00E+00 | 0.412 | 0.721 | GP |
| Gjc3 | -0.6260327 | 2.63E-04 | 1.00E+00 | 0.941 | 0.996 | GP |
| Slc22a23 | -0.6170532 | 6.15E-04 | 1.00E+00 | 0.588 | 0.783 | GP |
